# An atlas of microtubule lattice parameters regulated through ligand binding to the microtubule stabilizing sites

**DOI:** 10.1101/2025.11.11.687833

**Authors:** Daniel Lucena-Agell, Óscar Fernández, Rebeca París-Ogáyar, Juan Estévez-Gallego, Denisa Ondrúšková, Beatriz Álvarez-Bernad, Francesca Bonato, Jaime Larraga, Diego Ortíz de Elguea, Gregorio Javier Cano, Maarten Scheers, Hiroshi Imai, Toshiki Yagi, Hiroyuki Iwamoto, Juan Carlos Martínez-Guil, A. Jonathan Singh, Robert Alexander Keyzers, Christopher D. Vanderwal, Karl-Heinz Altmann, Johan Van der Eycken, Valle Palomo, Wei-Shuo Fang, Federico Gago, Zdeněk Lánský, Marcus Braun, María Á. Oliva, Shinji Kamimura, J. Fernando Díaz

**Author notes:** These authors have contributed equally (Daniel Lucena-Agell, Óscar Fernández, Rebeca París-Ogáyar).

## Abstract

Microtubules (MTs) are dynamic cytoskeletal polymers whose lattice architecture regulates force generation, nucleotide hydrolysis, and recognition by motor proteins and microtubule-associated proteins (MAPs). Microtubule-stabilizing agents (MSAs), including taxanes and laulimalide/peloruside-site ligands, suppress depolymerization by binding to defined lattice sites, yet stabilization is not structurally neutral. How ligand chemistry reshapes lattice organization and function remains unresolved.

Here, we address three mechanistic questions. First, do distinct ligand classes induce defined lattice states? Using X-ray fiber diffraction, we show that MSAs selectively stabilize two preferred longitudinal conformations, a compact state (∼4.06 nm monomer rise) and an expanded state (∼4.17 nm), while modulating lateral organization reflected in shifts in mean MT radius. These axial spacings cluster around discrete values across chemotypes, indicating stabilization of pre-existing conformational minima rather than continuous distortion. Second, are these states interconvertible upon changes in ligand occupancy? Time-resolved diffraction reveals that longitudinal transitions occur within seconds of ligand addition even at substoichiometric occupancy; whereas, lateral equilibration proceeds slower, consistent with redistribution within heterogeneous protofilament organizations. Third, do such structural states alter nucleotide hydrolysis and motor/MAP behavior? Expanded lattices are associated with reduced apparent GTP hydrolysis rates under steady-state assembly conditions and altered kinesin motility, whereas compact lattices preferentially promote tau binding and distinct motor interaction profiles.

Together, these findings establish longitudinal lattice conformation as a regulatory parameter and position microtubule-stabilizing agents as chemical tools that bias a dynamic structural landscape with predictable catalytic and transport consequences.

**Significance statement:** Microtubules generate force and support intracellular transport through lattice geometries selectively recognized by motor proteins and microtubule-associated proteins. Microtubule-stabilizing drugs such as taxanes and epothilones are widely used in chemotherapy but can cause neurotoxicity, likely because stabilization alters lattice architecture rather than simply preventing depolymerization. Here, we show that stabilizing ligands bias microtubules between two preferred longitudinal conformations, compact and expanded, that can switch within seconds of binding, while lateral organization adjusts slower. These structural states differentially regulate GTP hydrolysis and recognition by tau and kinesin, linking lattice geometry to catalytic and transport functions. By establishing lattice conformation as a tunable regulatory parameter, this work provides a framework for interpreting drug effects and designing structure-selective stabilizers with improved therapeutic profiles.

## Introduction

Microtubules (MTs) are dynamic cytoskeletal polymers essential for force generation, maintenance of cell shape, and polarized intracellular transport (1). They are assembled from αβ-tubulin heterodimers, whose conformational transitions couple nucleotide hydrolysis to polymerization, enabling mechanical work. Like actin, tubulin operates through an assembly state-dependent conformational change known as the cytomotive switch (2). In tubulin, this switch involves a curved-to-straight transition characterized by interdomain rotation and displacement of helix H7 (3, 4).

Because this conformational transition underlies controlled force generation, compounds that interfere with it disrupt processes such as cell division and angiogenesis. Many organisms have therefore evolved tubulin-targeting anticytomotive compounds (5), which bind to at least eight distinct sites on αβ-tubulin (6-8). Six of these sites are associated with microtubule-destabilizing agents (MDAs), including colchicine, vinblastine, maytansine, pironetin, gatorbulin, and todalam. These compounds block the cytomotive switch either by trapping tubulin in the curved conformation (e.g., colchicine, mebendazole and gatorbulin) or by disrupting longitudinal interdimer contacts (e.g., vinblastine, maytansine and pironetin), thereby preventing polymerization and collapsing the microtubule network (9-12). While highly effective against proliferating tumors, their mechanism of action involves the destruction of microtubules leading to one of the most limiting side effects in successful cancer therapy: peripheral neurotoxicity, caused by damage or degeneration of peripheral nerves (13).

Paclitaxel (Taxol®, PTX), approved in 1992 (14), inaugurated a second class of microtubule-targeting agents (MTAs): microtubule-stabilizing agents (MSAs). This group includes docetaxel (DTX), cabazitaxel (CBTX), and ixabepilone (IBP), which block the cytomotive switch in the straight-to-curved direction, thereby stabilizing polymerized MTs. Most MSAs [taxanes, epothilones, discodermolides (DDM), zampanolides (ZMP)] bind to the taxane site, reinforcing lateral contacts by stabilizing the M loop (15). In contrast, laulimalide (LAU)- and peloruside A (PLA)-site ligands stabilize MTs by bridging adjacent protofilaments (16). Although stabilization was initially thought to preserve structural roles while selectively affecting force generation, MSAs exhibit neurotoxicity comparable to destabilizing agents (17, 18).

Structural analyses reveal that stabilization is not structurally neutral. PTX and DTX induce longitudinal lattice expansion, increasing the monomer repeat from 4.06 nm to 4.17 nm and, in some cases, altering interprotofilament angles (15). In contrast, LAU-site ligands stabilize MTs without inducing axial expansion, though they may distort lateral interfaces. Notably, compounds such as pelophen B (PPH) can stabilize MTs without detectable lattice distortion (19). These observations suggest that stabilization and lattice geometry are separable properties.

MT structure also regulates its own biochemical activity. GTP hydrolysis at the β-tubulin exchangeable site, activated across the interdimer interface, establishes a nucleotide gradient along the lattice that is translated into subtle structural differences recognized by microtubule-associated proteins (MAPs) (1, 20). MAPs, including EB proteins, tau, and motors, selectively recognize specific lattice conformations. For example, kinesin-1 displays enhanced affinity for expanded lattices (21), whereas tau preferentially binds compacted lattices (22). These findings indicate that MAPs function as both sensors and modulators of lattice structure, and that drug-induced structural distortions are likely to alter MAP behavior (23).

Given the clinical relevance of tubulin-targeted therapies, it is essential to understand how MSAs modulate microtubule architecture and its functional consequences. Here, we systematically compare multiple chemotypes targeting both stabilizing sites under identical conditions. Using fiber diffraction of aligned MTs, molecular dynamics simulations, and TIRF-based single-molecule assays, we define discrete lattice states and examine their kinetic and functional implications for nucleotide hydrolysis and MAP interaction. This integrative framework provides a structural atlas of ligand-induced lattice states and offers tools to chemically manipulate microtubule architecture with precision.

## Results

### Distinct stabilizing-site ligands induce different lattice states

The objective of the present study was to determine how ligand chemistry modulates microtubule architecture and function. To address this, we focused on three mechanistic questions: (i) do distinct ligand classes induce defined lattice states; (ii) are these states interconvertible upon changes in ligand occupancy; and (iii) do such structural states alter nucleotide hydrolysis and motor/MAP behavior? The following sections address these questions using complementary structural, kinetic, and functional approaches.

Previous work from our group demonstrated that PTX (15, 24, 25) and other taxanes (15, 26, 27) markedly influence microtubule (MT) architecture, affecting both longitudinal lattice spacing and lateral organization. A recurring feature is lattice longitudinal expansion, reflected by an increase in monomer rise from ∼4.06 nm to ∼4.17 nm. Similar expanded conformations have been reported for hyper-stabilized MTs assembled with GMPCPP (28-30) and for catalytically inactive Gluα254A and Gluα254N MTs (31). However, expansion is not universally associated with activation. GMPPCP- and metal fluoride (BeF_3_^−^ and AlF_4_^−^)-stabilized MTs can adopt compressed states (25), and PTX- and BeF_3_^−^-bound MTs likewise stabilize a compressed lattice (25). Conversely, expansion can be induced by BIII, a PTX precursor lacking the C-13 side chain and displaying minimal activity (15). Together, these observations indicate that longitudinal lattice expansion reflects specific structural effects of ligand binding rather than tubulin activation *per se*.

To systematically define how ligand chemistry regulates MT structure, we analyzed aligned MTs by X-ray fiber diffraction (32) in the presence of a broad panel of MT-stabilizing agents (MSAs) targeting either the taxane or the peloruside/laulimalide sites (**Figure 1**). This library comprises taxanes, epothilones, DDM, ZMPs, cyclostreptin (CSTR), taccalonolide AJ (TACCA-AJ), LAU and pelorusides. Previously reported datasets (15, 19, 25, 33) were re-measured or re-analyzed under identical conditions to ensure strict internal consistency across the entire dataset (**Table I**). Additional conformational data obtained in the presence of γ-phosphate or nucleotide analogs are presented in **Table SI**.

**Figure 1.**
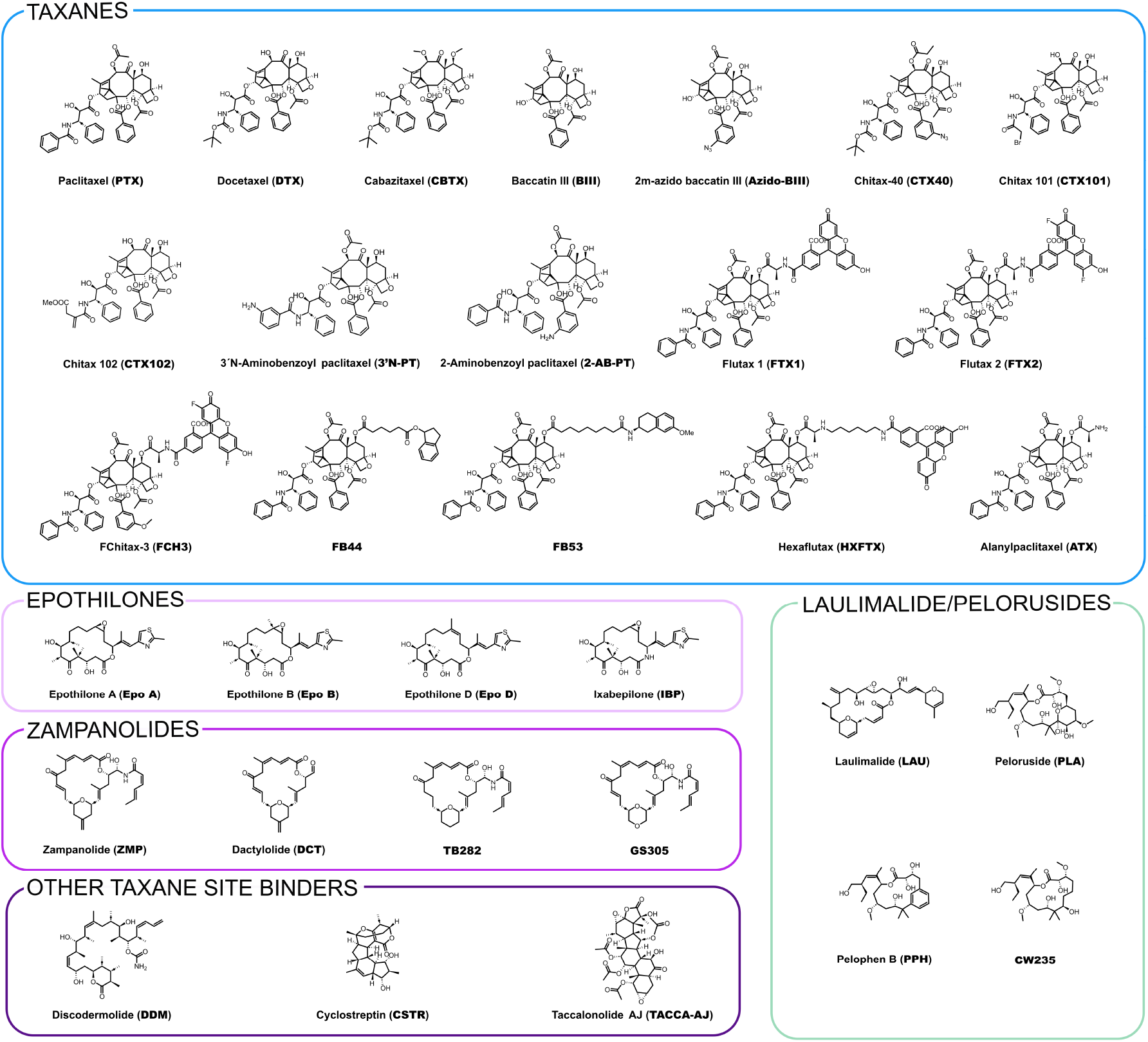
Chemical structures of the microtubule-stabilizing ligands used in this study, grouped by chemotype and binding site (taxane-site ligands and peloruside/laulimalide-site ligands).

**Table I.**
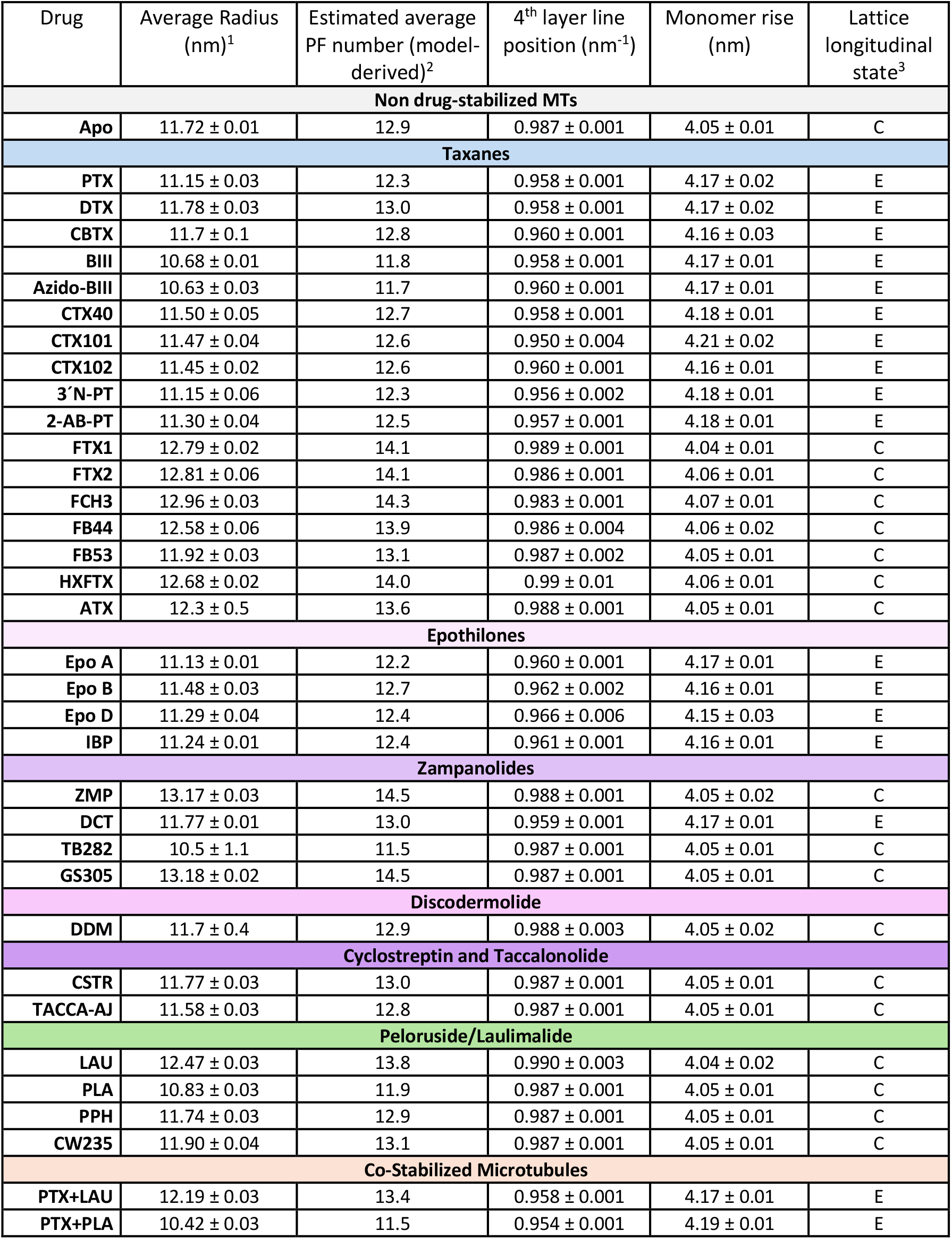
Structural parameters of microtubules in the presence of stabilizing ligands. 1.- Reported SEM values correspond to the uncertainty in the determination of the mean microtubule radius from diffraction measurements and do not reflect the dispersion of protofilament numbers within the heterogeneous microtubule population. 2.- Protofilament (PF) numbers were estimated from the measured mean MT radius using a geometric conversion [assuming an average inter-protofilament distance of 5.7 nm (26)] and are provided for comparative purposes only. Throughout the manuscript, longitudinal spacing and mean radius are treated as primary observables, whereas PF number serves as a derived descriptor of lateral organization within heterogeneous MT populations. Protofilament numbers are therefore rounded to the nearest decimal. 3.- Lattice longitudinal state classification: C, compressed lattice; E, expanded lattice.

| Drug | Average Radius (nm)<sup>1</sup> | Estimated average PF number (model-derived)<sup>2</sup> | 4<sup>th</sup> layer line position (nm<sup>-1</sup>) | Monomer rise (nm) | Lattice longitudinal state<sup>3</sup> |
| --- | --- | --- | --- | --- | --- |
| **Non drug-stabilized MTs** |  |  |  |  |  |
| **Apo** | 11.72 ± 0.01 | 12.9 | 0.987 ± 0.001 | 4.05 ± 0.01 | C |
| **Taxanes** |  |  |  |  |  |
| **PTX** | 11.15 ± 0.03 | 12.3 | 0.958 ± 0.001 | 4.17 ± 0.02 | E |
| **DTX** | 11.78 ± 0.03 | 13.0 | 0.958 ± 0.001 | 4.17 ± 0.02 | E |
| **CBTX** | 11.7 ± 0.1 | 12.8 | 0.960 ± 0.001 | 4.16 ± 0.03 | E |
| **BIII** | 10.68 ± 0.01 | 11.8 | 0.958 ± 0.001 | 4.17 ± 0.01 | E |
| **Azido-BIII** | 10.63 ± 0.03 | 11.7 | 0.960 ± 0.001 | 4.17 ± 0.01 | E |
| **CTX40** | 11.50 ± 0.05 | 12.7 | 0.958 ± 0.001 | 4.18 ± 0.01 | E |
| **CTX101** | 11.47 ± 0.04 | 12.6 | 0.950 ± 0.004 | 4.21 ± 0.02 | E |
| **CTX102** | 11.45 ± 0.02 | 12.6 | 0.960 ± 0.001 | 4.16 ± 0.01 | E |
| **3'-N-PT** | 11.15 ± 0.06 | 12.3 | 0.956 ± 0.002 | 4.18 ± 0.01 | E |
| **2-AB-PT** | 11.30 ± 0.04 | 12.5 | 0.957 ± 0.001 | 4.18 ± 0.01 | E |
| **FTX1** | 12.79 ± 0.02 | 14.1 | 0.989 ± 0.001 | 4.04 ± 0.01 | C |
| **FTX2** | 12.81 ± 0.06 | 14.1 | 0.986 ± 0.001 | 4.06 ± 0.01 | C |
| **FCH3** | 12.96 ± 0.03 | 14.3 | 0.983 ± 0.001 | 4.07 ± 0.01 | C |
| **FB44** | 12.58 ± 0.06 | 13.9 | 0.986 ± 0.004 | 4.06 ± 0.02 | C |
| **FB53** | 11.92 ± 0.03 | 13.1 | 0.987 ± 0.002 | 4.05 ± 0.01 | C |
| **HXFTX** | 12.68 ± 0.02 | 14.0 | 0.99 ± 0.01 | 4.06 ± 0.01 | C |
| **ATX** | 12.3 ± 0.5 | 13.6 | 0.988 ± 0.001 | 4.05 ± 0.01 | C |
| **Epothilones** |  |  |  |  |  |
| **Epo A** | 11.13 ± 0.01 | 12.2 | 0.960 ± 0.001 | 4.17 ± 0.01 | E |
| **Epo B** | 11.48 ± 0.03 | 12.7 | 0.962 ± 0.002 | 4.16 ± 0.01 | E |
| **Epo D** | 11.29 ± 0.04 | 12.4 | 0.966 ± 0.006 | 4.15 ± 0.03 | E |
| **IBP** | 11.24 ± 0.01 | 12.4 | 0.961 ± 0.001 | 4.16 ± 0.01 | E |
| **Zampanolides** |  |  |  |  |  |
| **ZMP** | 13.17 ± 0.03 | 14.5 | 0.988 ± 0.001 | 4.05 ± 0.02 | C |
| **DCT** | 11.77 ± 0.01 | 13.0 | 0.959 ± 0.001 | 4.17 ± 0.01 | E |
| **TB282** | 10.5 ± 1.1 | 11.5 | 0.987 ± 0.001 | 4.05 ± 0.01 | C |
| **GS305** | 13.18 ± 0.02 | 14.5 | 0.987 ± 0.001 | 4.05 ± 0.01 | C |
| **Discodermolide** |  |  |  |  |  |
| **DDM** | 11.7 ± 0.4 | 12.9 | 0.988 ± 0.003 | 4.05 ± 0.02 | C |
| **Cyclostreptin and Taccalonolide** |  |  |  |  |  |
| **CSTR** | 11.77 ± 0.03 | 13.0 | 0.987 ± 0.001 | 4.05 ± 0.01 | C |
| **TACCA-AJ** | 11.58 ± 0.03 | 12.8 | 0.987 ± 0.001 | 4.05 ± 0.01 | C |
| **Peloruside/Laulimalide** |  |  |  |  |  |
| **LAU** | 12.47 ± 0.03 | 13.8 | 0.990 ± 0.003 | 4.04 ± 0.02 | C |
| **PLA** | 10.83 ± 0.03 | 11.9 | 0.987 ± 0.001 | 4.05 ± 0.01 | C |
| **PPH** | 11.74 ± 0.03 | 12.9 | 0.987 ± 0.001 | 4.05 ± 0.01 | C |
| **CW235** | 11.90 ± 0.04 | 13.1 | 0.987 ± 0.001 | 4.05 ± 0.01 | C |
| **Co-Stabilized Microtubules** |  |  |  |  |  |
| **PTX+LAU** | 12.19 ± 0.03 | 13.4 | 0.958 ± 0.001 | 4.17 ± 0.01 | E |
| **PTX+PLA** | 10.42 ± 0.03 | 11.5 | 0.954 ± 0.001 | 4.19 ± 0.01 | E |

Fiber diffraction patterns revealed two principal structural effects of ligand binding (**Figure 2**). First, several ligands induce a clear shift in longitudinal lattice spacing, consistent with previous observations (25, 34). This shift is visible as an inward displacement of the meridional layer lines arising from the helical repeat (located at 1/4, 2/4, 3/4 and 4/4 nm^−1^ in the diffractogram; see Materials and Methods) (**Figures 2C–F**), corresponding to an increase in monomer rise from ∼4.06 nm (compressed state) to ∼4.17 nm (expanded state). The expansion is asymmetric, and a reflection at ∼1/8 nm^−1^ becomes detectable, consistent with the emergence of the full ∼8 nm axial periodicity of the αβ-tubulin dimer. Across the analyzed ligands, the longitudinal spacing clustered around these two preferred values, consistent with two interconvertible longitudinal lattice states rather than broad continuous variation. Notably, under some co-stabilization conditions, specifically GMPCPP-stabilized microtubules bound to ligands that favor compression, intermediate or broadened layer lines were observed, indicative of structural heterogeneity within the MT population (**Table SI**).

**Figure 2.**
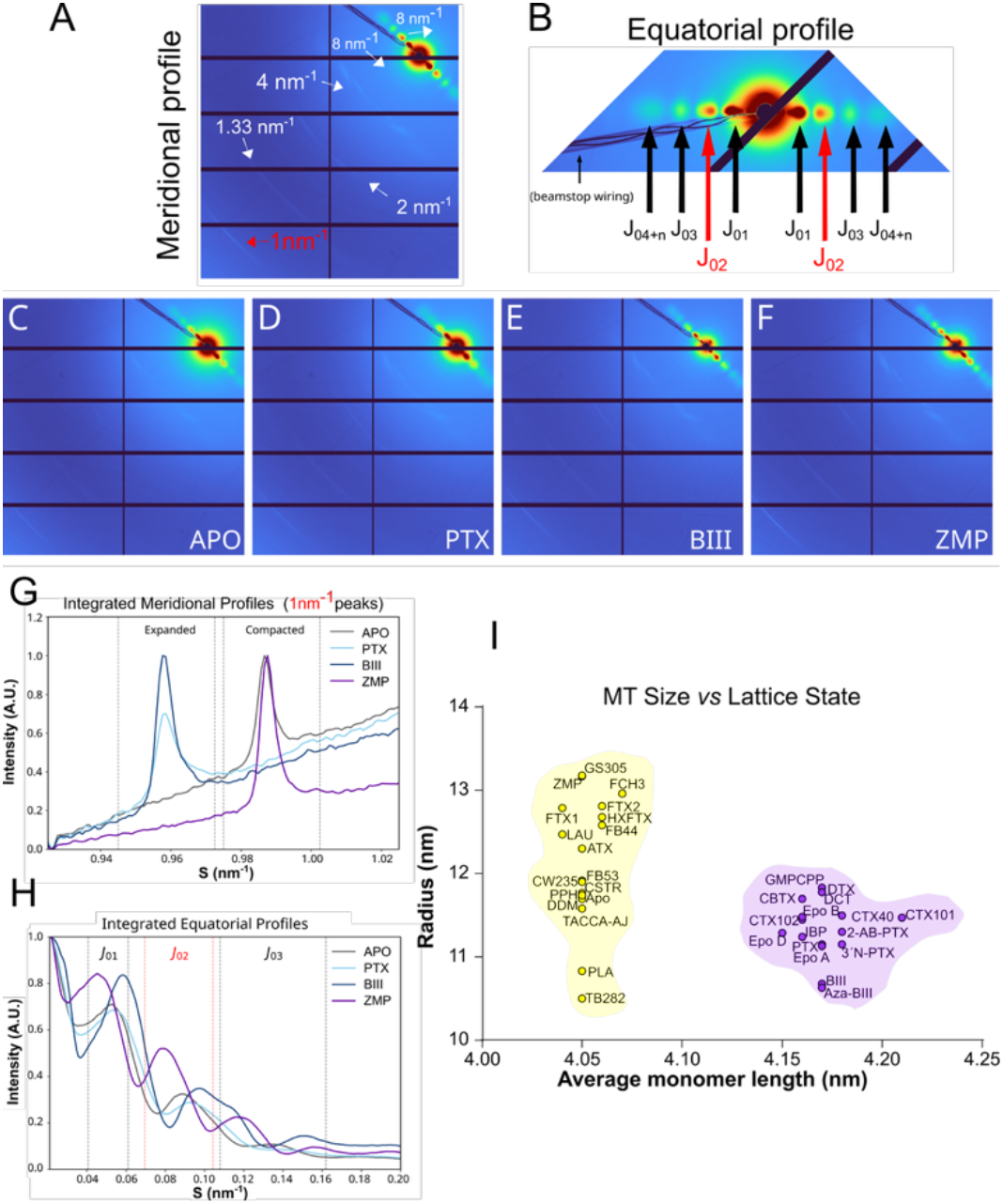
Interpretation of microtubule diffractograms. **A**. Reference image illustrating meridional signal interpretation and layer-lines localizations. **B**. Reference image illustrating equatorial signal interpretation and Bessel peak positions. **C–F**. Representative diffractograms for four microtubule conditions: (C) APO, (D) PTX, (E) BIII, and (F) ZMP. **G**. Azimuthally integrated meridional profiles focused on the 1 nm^-1^ layer line, highlighting the ranges associated with compacted and expanded lattice states. **H**. Azimuthally integrated equatorial profiles from the diffractograms above, including theoretical ranges for Bessel J_0_ peak positions corresponding to microtubules with radii between 10 and 15 nm. **I. Relationship between the MT lattice expansion state and MT radius for all the measured conditions**.

The axial spacing measured by fiber diffraction (∼40–42 Å) corresponds to half of the αβ-tubulin dimer repeat and is therefore consistent with the ∼81–84 Å dimer rise reported by cryo-EM. The apparent discrepancy arises because α- and β-tubulin are not individually resolved at the resolution of the diffraction experiment, whereas they are distinguished in cryo-EM reconstructions. Accordingly, the ∼1/4 nm^−1^ meridional reflection reports the monomer repeat, while the full dimer periodicity appears at ∼1/8 nm^−1^. This latter reflection is prominent in expanded lattices but weak or absent in compressed lattices (**Figures 2C-F**), indicating that the longitudinal conformational change is largely asymmetric within the αβ-tubulin dimer.

Second, ligand binding modulates lateral lattice geometry, reflected through shifts in the position of the equatorial maxima of the diffractograms (**Figure 2G**). These changes correspond to variations in mean MT radius and lateral inter-protofilament organization within heterogeneous *in vitro*–assembled populations that contain a distribution of protofilament numbers. This is consistent with previous observations for both the taxane site (24, 26) and the peloruside/laulimalide site (19).

A consistent relationship between longitudinal and lateral parameters is observed: ligands that stabilize expanded lattices tend to maintain or slightly reduce the mean MT radius; whereas, compounds that preserve compressed axial spacing are frequently associated with increased mean radius. For example, BIII stabilizes an expanded lattice while reducing mean radius, while ZMP stabilizes a compressed lattice with increased mean radius (**Figure 2 and Table I**). Combinations of drugs with GMPCPP further illustrate this decoupling, resulting in MT populations with increased mean radius while maintaining an expanded longitudinal lattice (**Table SI**).

Importantly, not all taxane-site ligands induce lattice expansion. While most taxanes, except those modified at C-7 (33), and epothilones promote expansion, DDM, ZMP, CSTR, TACCA-AJ, and ligands targeting the peloruside/laulimalide site stabilize MTs without altering axial spacing, in agreement with previous observations (19). These findings indicate that stabilization through the taxane site can be mechanistically separable from longitudinal lattice expansion. Collectively, these results address the first mechanistic question by showing that ligand binding stabilizes defined lattice states, characterized by two preferred longitudinal conformations and ligand-dependent lateral organization. Notably, lattice state depends on specific chemical features rather than binding site alone, as closely related taxanes (including C-7-modified analogues) stabilize distinct conformations.

### Stoichiometry of the taxane site regulation of the microtubule structure

Having established that occupancy of the taxane site alters longitudinal lattice spacing and lateral geometry, we next examined the ligand:tubulin stoichiometry required to induce these structural transitions (**Figure 3**). This analysis provides insight into how partial site occupancy, more relevant to cellular conditions, modulates lattice organization.

**Figure 3.**
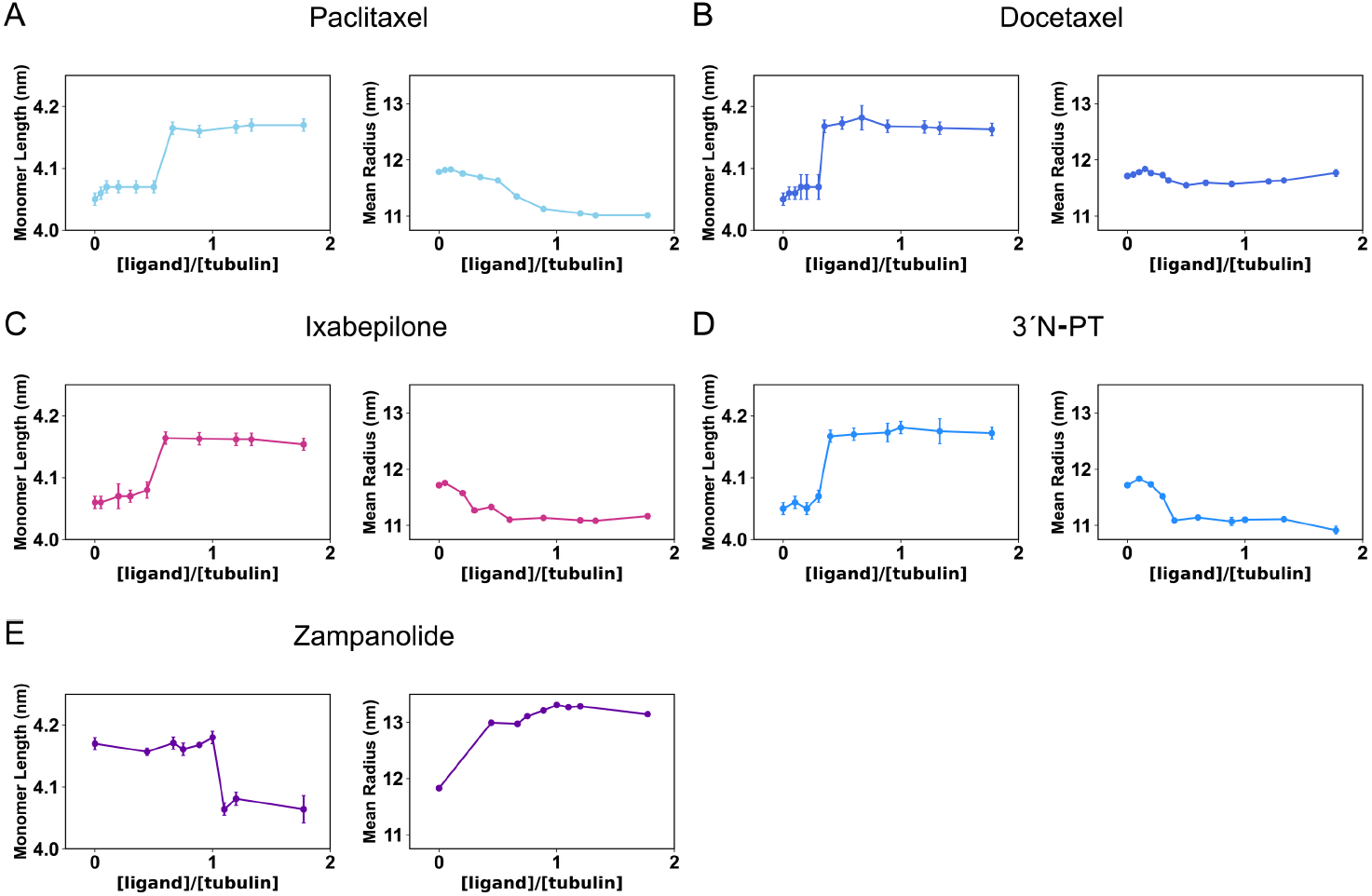
Structural response of apo- (compressed A, B, C, D) and GMPCPP- (expanded, E) microtubules to different ligand:tubulin ratios. The plots show the variation in average monomer length and mean radius, highlighting the impact of ligand concentration on lattice organization. (A) PTX, (B) DTX, (C) IBP, (D) 3’N-PT, (E) ZMP.

Five taxane-site ligands were selected: clinically used compounds PTX (**Figure 3A**), DTX (**Figure 3B**), and IBP (**Figure 3C**), a minimally modified fluorescent taxane derivative (3′N-PT) (**Figure 3D**), and the covalent binder ZMP (**Figure 3E**). PTX, IBP, and 3′N-PT induce lattice expansion and modify the mean radius, whereas DTX primarily modulates longitudinal spacing. By contrast, ZMP, which stabilizes a compressed lattice, was tested on pre-expanded GMPCPP-bound MTs.

Remarkably, lattice expansion was observed at substoichiometric ligand:tubulin ratios. For all expanding compounds tested, the longitudinal spacing approached that of the fully ligand-bound state at partial occupancy, typically around a 1:3–1:2 ratio. In contrast, ZMP induced re-compression of GMPCPP-expanded MTs but required near-stoichiometric occupancy to achieve this effect.

Importantly, the longitudinal response did not vary continuously with ligand concentration but clustered around the compressed (∼4.06 nm) or expanded (∼4.17 nm) states. Intermediate axial spacings were not resolved within experimental uncertainty. In contrast to the quasi-binary longitudinal transition, lateral organization is distributed across multiple protofilament architectures already present in the population. As drug concentration increases, the mean radius can therefore shift progressively through redistribution among these lateral states, producing a smoother ensemble transition. These findings indicate that partial occupancy of the taxane site is sufficient to bias the MT lattice toward defined structural states, consistent with a cooperative structural response of the polymer.

### Time-resolved fiber diffraction analysis of the structural effect of chemical stabilization of microtubules induced by MSAs

Having defined the stoichiometry required to induce structural transitions, next we examined how pre-formed MT lattices respond to acute changes in chemical environment (**Supp. Movie 1**). We analyzed three taxane-site ligands: DTX (minimal effect on mean radius), PTX, and BIII (both associated with reduced mean radius at equilibrium) - all of which stabilize expanded lattices - and compared them with laulimalide, which increases mean radius without altering axial spacing (19). Previous time-resolved measurements of non-aligned MTs showed that diameter changes following PTX or DTX addition occur within the ∼30 s dead time of the technique (35). Using a shear-flow mixing system with ∼1 s resolution, we independently monitored axial and radial parameters (**Figure 4**).

**Figure 4.**
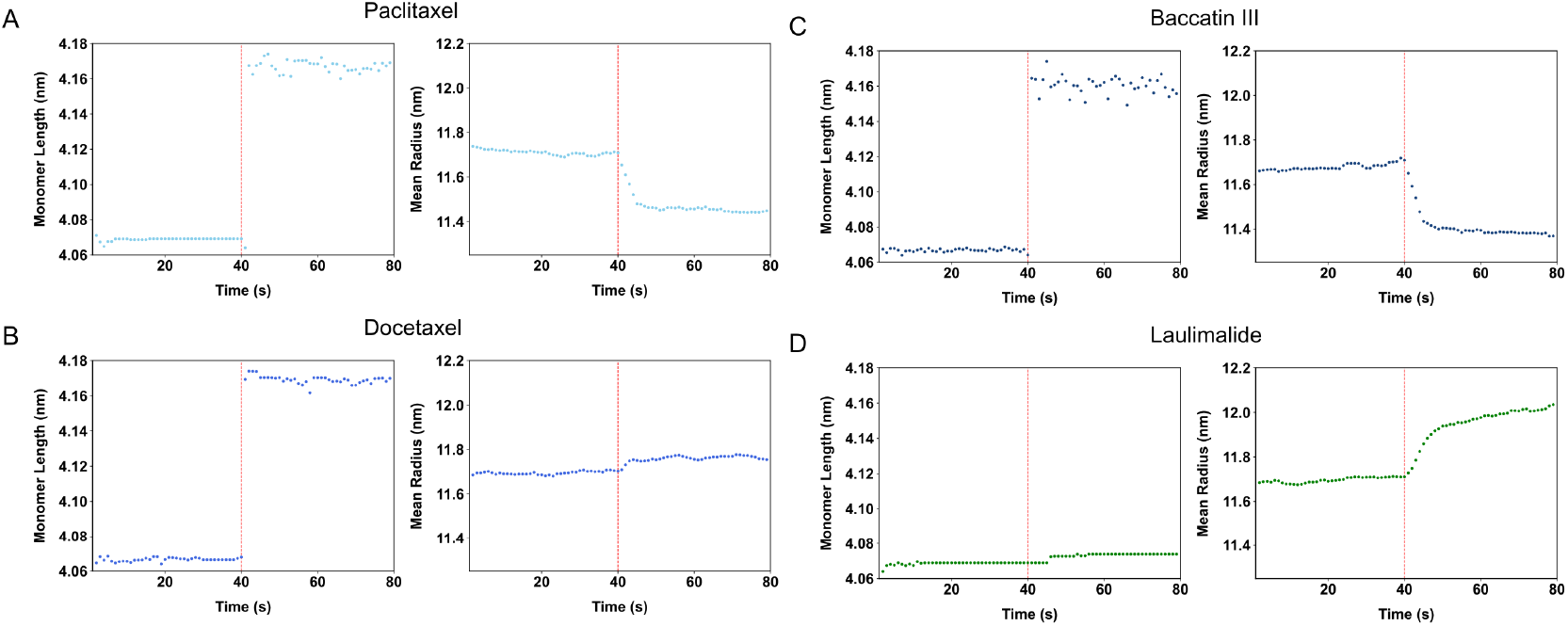
Time-resolved axial and radial measurements reveal different dynamics of microtubule lattice reorganization after ligand addition at 40 s (red dashed line). Left: average axial tubulin repeat. Right: average mean radius. Ligands: (A) PTX, (B) DTX, (C) BIII, (D) LAU.

Upon addition of PTX, DTX, or BIII, axial lattice expansion was detected within the first second of measurement. In all cases, the monomer rise shifted rapidly toward the expanded value (∼4.17 nm). Because this axial parameter is determined directly from meridional peak positions and does not rely on geometric modeling, these observations indicate that longitudinal lattice rearrangement occurs on a sub-second timescale. In contrast, addition of LAU, which does not alter axial spacing, produced no detectable change in monomer rise (**Figure 4D**), serving as an internal control.

The radial response differed markedly. Although equilibrium measurements indicate substantial differences in mean radius between apo- and drug-stabilized MTs (**Table I**), early time-resolved changes in mean radius were significantly smaller. For example, PTX-stabilized MTs assembled in the continuous presence of the drug exhibit, on average, ∼0.6 nm smaller radius than apo MTs (**Table I**), yet only a ∼0.3 nm decrease was measured within the first 3 s following addition to pre-assembled MTs. Similarly, BIII, associated with a larger equilibrium reduction (∼1.0 nm), produced only a modest early shift. Conversely, LAU produced a modest early increase (∼0.3 nm) in mean radius, consistent with its known equilibrium effect (19). Control experiments in which apo-assembled microtubules were incubated for 30 min with superstoichiometric concentrations of PTX, DTX, or BIII showed only a partial shift in mean radius toward the values of microtubules assembled in the presence of the drug (11.28 nm vs 11.15 nm for PTX; 11.02 nm vs 10.68 nm for BIII) (**Table S2**), indicating that full equilibration of lateral organization requires substantially longer times.

Importantly, the magnitude of the rapid radial change was consistently smaller than the equilibrium difference observed under steady-state conditions (**Table S2**). These results indicate that early structural responses primarily reflect rapid longitudinal lattice rearrangement, whereas changes in mean radius evolve more gradually. Given the heterogeneous nature of in vitro– assembled MT populations, the rapid radial shifts observed over time are consistent with fast incorporation of the available free tubulin pool (approximately 15%), rather than synchronous protofilament addition or loss.

The rapid and concentration-dependent transition of the longitudinal parameter toward the expanded state, even at partial occupancy, is consistent with a cooperative structural response of the polymer lattice, in which ligand binding at a subset of sites biases the conformational equilibrium of neighboring tubulin subunits. In contrast, the slower evolution of mean radius suggests that lateral reorganization to the equilibrium state proceeds through progressive redistribution within the existing protofilament population rather than an immediate, concerted structural rearrangement.

Together, these findings address the second mechanistic question by demonstrating that ligand-induced lattice states are interconvertible, rapidly in the longitudinal dimension, whereas lateral equilibration toward the final structural distribution occurs on a longer timescale.

### Molecular modeling and dynamics of ligand-modulated lattice states

To gain structural insight into how taxane-site ligands modulate lattice parameters and influence catalytic geometry, we performed normal mode analysis and molecular dynamics (MD) simulations on representative compacted and expanded microtubule interfaces. We examined (i) low-frequency collective motions in 13- and 14-protofilament lattice fragments, (ii) the conformational behavior of β1:α2 interdimer interfaces in apo and drug-bound systems (DTX, Epo A, ZMP, DDM, and ATX), and (iii) minimal three-protofilament lattice models as previously described (36). Representative compacted and expanded MT turns have been deposited in the ModelArchive repository (37) (codes listed in Materials and Methods).

Normal mode analysis consistently identified a dominant low-frequency collective motion corresponding to cooperative bending and longitudinal closure of the lattice (**Supp. Movie 2**). Sampling along this mode generated conformations consistent with subtle variations in axial rise and curvature, supporting the experimentally observed sensitivity of longitudinal spacing and mean radius to small structural fluctuations.

Comparison of β1:α2 interfaces in GTP-versus GDP-bound states reproduced the expected differences in γ-phosphate coordination and hydrogen-bonding networks at the E-site. The simulations revealed that the interdimer interface is structurally plastic and capable of sampling closely related conformational states distinguished by coordinated rearrangements of side chains, nucleotide phosphates, and associated water molecules (**Figure S2**).

When ligands were included at the taxane site, compacted and expanded lattices exhibited distinct interdimer rise distances (approximately 40.5 Å and 42.0 Å, respectively), in agreement with diffraction measurements. Ligand binding altered the conformational ensemble of the β1:α2 interface and modified the dynamic coupling between the taxane pocket and the nucleotide-binding interface. These changes were accompanied by subtle rearrangements of the hydrogen-bond network and phosphate geometry at the E-site, consistent with the experimentally observed modulation of GTP hydrolysis.

Together, these simulations show that ligand binding stabilizes distinct conformational ensembles of the interdimer interface, linking longitudinal lattice spacing to local remodeling of catalytic-site geometry.

### Effect of microtubule structure on nucleotide hydrolysis

Because ligand binding modulates longitudinal lattice spacing, we examined whether defined lattice states are associated with differences in GTP hydrolysis at the E-site of β-tubulin, which becomes catalytically activated upon incorporation into the microtubule lattice through Gluα254 of the incoming α-tubulin subunit (38).

First, we verified that drug-stabilized microtubules retain GTPase activity by analyzing nucleotide content in assembled polymers. After one hour of incubation, all drug-bound MTs contained approximately one mole of GTP and one mole of GDP per tubulin dimer, consistent with continued GTP hydrolysis under the experimental conditions.

We next quantified apparent GTP hydrolysis rates under steady-state assembly conditions of MTs in the presence of expanding ligands (PTX, Epo A) and non-expanding ligands bound to either the taxane site (DDM, ZMP) or the peloruside/laulimalide site (PPH) (19), relative to apo MTs, using the malachite green assay (**Table II**). As described in the Materials and Methods (see **SI**), phosphate accumulation was measured after acid quenching and therefore reflects cumulative GTP hydrolysis associated with polymerized tubulin. Because the reaction was quenched under strongly denaturing conditions, the assay quantifies total inorganic phosphate generated at the time of sampling rather than phosphate release kinetics. GTPase rates were determined from the linear regime of phosphate accumulation (**Figure S1**) (early time points corresponding to nucleation and rapid elongation were excluded), a phase in which polymer mass is at steady-state and phosphate concentration increases linearly with time. The slope of this region (normalized by tubulin concentration) was taken as the apparent rate.

**Table II.**
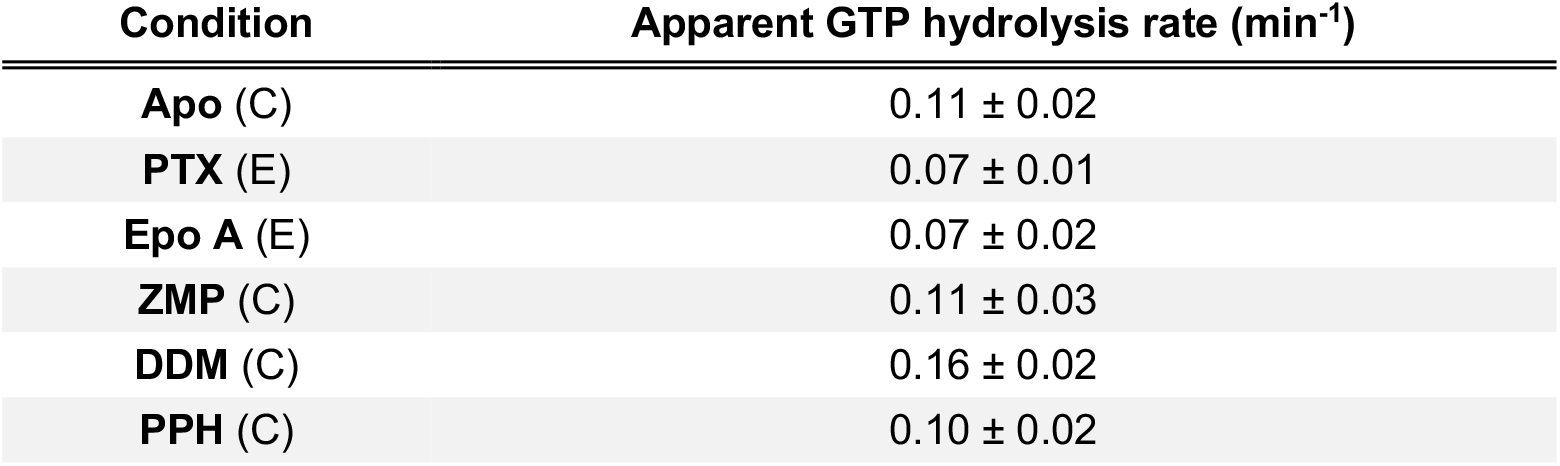
Apparent GTP hydrolysis rate of drug-stabilized expanded and compressed microtubules. (C) compressed and (E) expanded state of tubulin.

Expanded lattices stabilized by PTX or Epo A exhibited reduced hydrolysis rates relative to apo MTs (∼35% decrease). In contrast, compressed lattices stabilized by ZMP or PPH displayed rates similar to apo MTs, while DDM-bound MTs showed increased hydrolysis (∼45% increase). Thus, modulation of longitudinal lattice state is associated with differences in apparent GTPase activity under steady-state assembly conditions.

Because E-site catalysis is highly sensitive to the geometric arrangement of catalytic residues, nucleotide phosphates, and associated water molecules (39), these differences are consistent with subtle structural changes propagated from the ligand-binding pocket to the catalytic interface.

### Effect of structural changes on kinesin-driven intracellular transport on microtubules

Kinesin-driven intracellular transport along microtubules, essential for neurotransmitter trafficking in axons, is a major cellular function of the microtubule cytoskeleton (40). This process depends on mechanochemical coupling between kinesin stepping dynamics and recognition of the microtubule lattice. Having established tools to modulate microtubule structure, we next examined how defined lattice states affect kinesin-1 motility.

To address this question, we employed the time-lapse total internal reflection fluorescence (TIRF) microscopy system described previously (33). Using fluorescently labeled kinesin motors (Kif5B 1–905–eGFP (41)), we tracked individual motors moving along immobilized MTs stabilized by a panel of distinct MSAs that generate different lattice architectures (**Figure 5 and Supp. Movie 3**). We quantified landing rate (events·s^-1^·μm^-1^ of MT·nM^-1^ kinesin), run duration, run length, and velocity.

**Figure 5.**
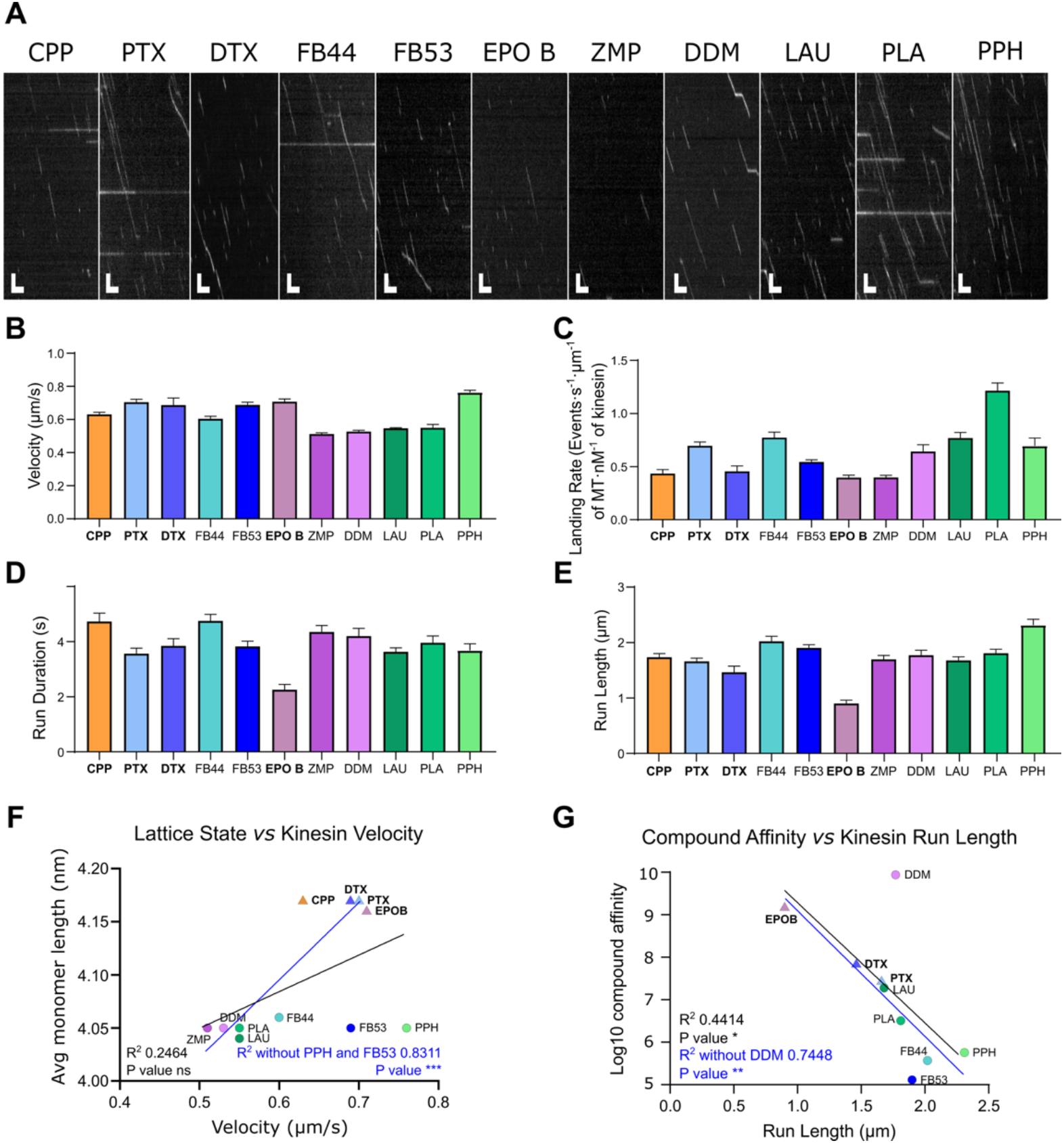
Characterization of kinesin motility. **A**. Representative kymographs of single Kif5B-eGFP motors moving along a stabilized microtubule. Vertical scale bar: 2 µm; horizontal scale bar: 10 s. **B–E**. Bar plots and quantitative summary from at least three independent experiments, showing (B) mean velocity (µm/s), (C) mean landing rate of kinesin (events·s^−1^·μm^−1^ of MT·nM^−1^ kinesin), (D) mean run duration (s), and (E) mean run length (µm), of single Kif5B-eGFP molecules moving on stabilized MTs. Error bars represent the SEM. Data were pooled from ≥ 2 experiments per condition, analyzing between 36–48 MTs per replica. **F-G**. Statistical correlations found between the structural and affinity parameters of the drug-stabilized lattices and the interaction of kinesin with them. Pearson correlation was obtained using Sigmaplot 16.0 software to determine R square and P value (**** p < 0.0001; *** p < 0.001, ** p < 0.01 and * p < 0.05). Compound binding affinities were taken from (19, 33, 42-44).

Data for PTX (expanded), FTX2 (compressed), FB44 (compressed), FB53 (compressed), and GMPCPP (expanded) were taken from our earlier work (33). We extended the analysis to additional taxane-site ligands, DTX (expanded), Epo B (expanded), ZMP (compressed), and DDM (compressed), as well as peloruside/laulimalide-site ligands: PLA, LAU, and PPH (all compressed).

Although kinesin-MT interactions show compound-specific complexity, several consistent trends emerge. Kinesin velocity shows a clear dependence on longitudinal lattice state (**Figures 5B, 5F**), with higher velocities observed on expanded microtubules. In contrast, compressed lattices display a broader distribution of velocities. Within this group, slower velocities are generally associated with higher-affinity ligands (e.g., ZMP, DDM, LAU, PLA) (19, 42-44), whereas compressed lattices stabilized by lower-affinity ligands (e.g., FB44, FB53, PPH) (19, 33) support faster movement. Thus, while expanded lattices consistently support higher kinesin velocities, compaction alone does not determine stepping speed, indicating that ligand-specific effects modulate motor dynamics within compressed lattices.

Landing rate (**Figure 5C**) is reduced on expanded lattices and in the presence of ZMP. Run duration (**Figure 5D**) is largely similar across conditions, with the exception of Epo B, which produces significantly shorter dwell times. Run length (**Figure 5E**) remains relatively stable, with two notable deviations: Epo B reduces run length, whereas PPH increases it. Interestingly, run length shows a partial correlation with ligand binding affinity (**Figure 5G**). Given that all ligands were tested at a fixed concentration (10 μM), this relationship should be interpreted cautiously, as binding affinity may influence both lattice stabilization and fractional occupancy. However, the trend persists across compounds with similar expected occupancy, suggesting that it reflects intrinsic ligand-dependent effects on lattice flexibility and motor interaction rather than differences in binding extent.

Epo B is notable in that it strongly stabilizes an expanded lattice (K_b_ = 1.5 × 10^9^ M^−1^ at 26 °C) (42). Despite maintaining high stepping velocity, kinesin exhibits reduced interaction time and shorter run length on Epo B-stabilized lattices, indicating that lattice expansion alone does not determine motor processivity.

### Effect of structural changes on tau binding and kinetics

Tau is the principal microtubule-associated protein in axons and requires dynamic interaction with microtubules (MTs) to exert its stabilizing and protective function (45). Impaired tau-MT interaction contributes to neurofibrillary tangle formation in Alzheimer’s disease and compromises axonal MT stability. To determine how specific lattice states influence tau recognition, we analyzed the binding of mCherry-labeled tau (441 isoform) to stabilized MTs using the same TIRF microscopy platform employed for kinesin assays (33).

Distinct ligand-stabilized MTs were immobilized in flow chambers and visualized by TIRF microscopy (**Figure 6A**). Tau–mCherry binding was examined at 5 nM, 10 nM, and 40 nM (**Figure 6B–C**). ZMP, DDM, PLA, and LAU were analyzed at all three concentrations, whereas the remaining ligands were tested at 10 and 40 nM. Binding was monitored over time, and both spatial coverage along MT length and integrated fluorescence intensity per micron were quantified.

**Figure 6.**
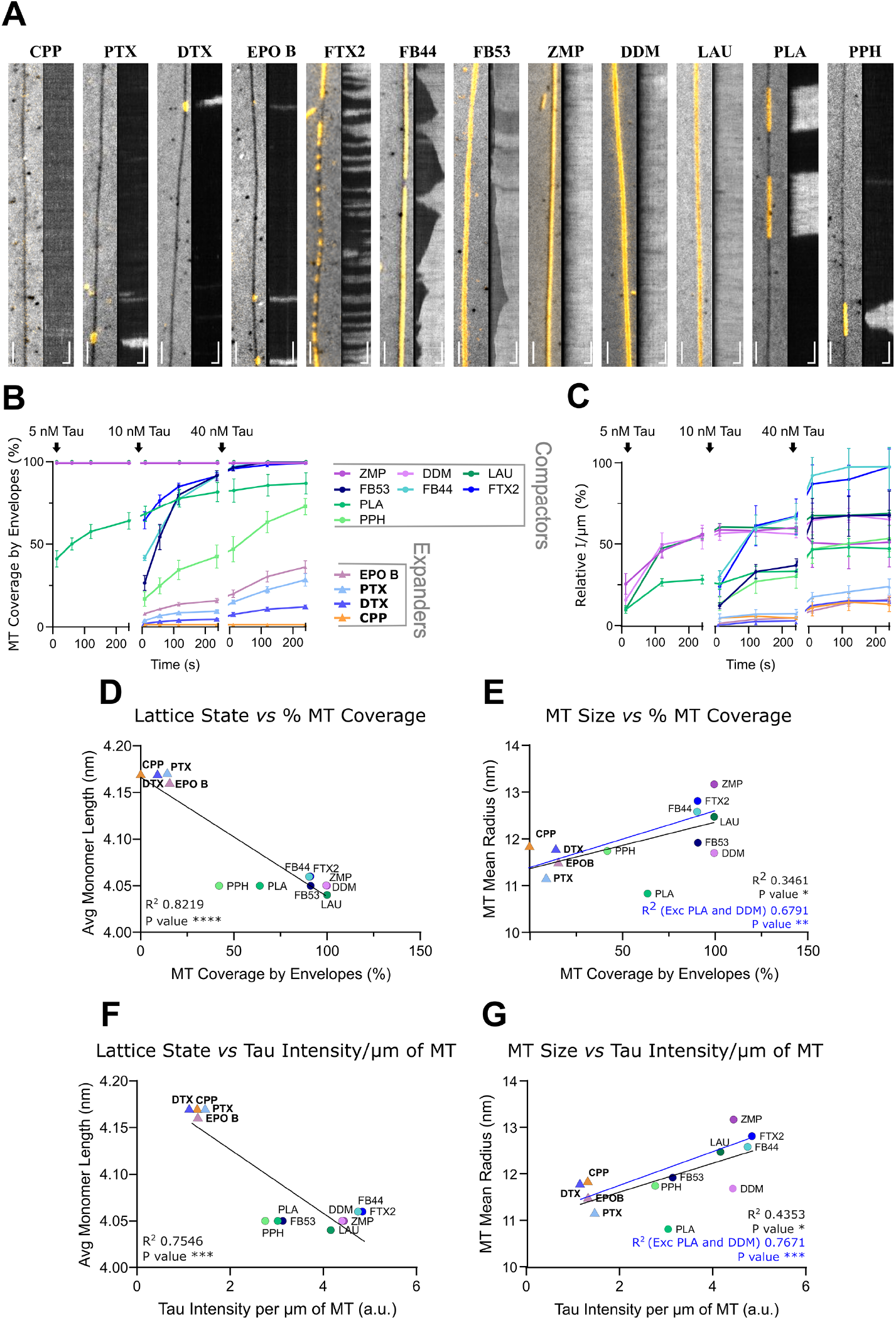
Characterization of tau interaction with microtubules. **A**. Representative fluorescence images (after 4 minutes) and corresponding kymographs of tau-mCherry binding to MTs stabilized with GMPCPP or different drugs. Vertical scale bars: 2 µm; horizontal scale bars: 60 s. Taxol derivatives are shown in blue shades, other taxanes in purple tones, and peloruside/laulimalide site ligands in different greens. Triangle symbols indicate expanders; circle symbols represent non-expander compounds. **B**. Fraction of MT length covered by tau envelopes over time following tau-mCherry addition. **C**. Integrated fluorescence density per µm of MT over time after tau-mCherry addition. **D-G**. Statistical correlations found between the structural parameters of the drug stabilized lattices and the binding of tau to them. Pearson correlation was obtained using Sigmaplot 16.0 software to determine R square and P value (**** p < 0.0001; *** p < 0.001, ** p < 0.01 and * p < 0.05).

Tau binding patterns correlated strongly with longitudinal lattice state (**Figures 6D** and **6F**), and four distinct recognition behaviors were observed. First, compact lattices stabilized by ZMP, DDM, and LAU were recognized extremely rapidly and became homogeneously coated along their entire length (**Figure 6B; Supp. Movies 4–6**). Even at 5 nM tau, near-complete spatial coverage was reached within ∼10 s (the dead time of the assay), without detectable patch formation. Although total fluorescence continued to increase over time, spatial coverage was essentially immediate, indicating rapid uniform recognition followed by progressive occupancy. Second, other compact lattices (PLA, PPH, FB44, FB53, FTX2) also supported rapid tau binding (**Figure 6B**) but through a spatially heterogeneous mechanism. Binding initiated in high-density patches that grow progressively along the MT (**Supp. Movies 7–8**), consistent with cooperative envelope formation (46, 47). Recognition was therefore favorable, but spatial propagation required local reorganization. Third, GMPCPP-stabilized MTs, constrained in an expanded conformation, exhibited negligible tau binding under these conditions (**Figure 6B**), indicating that this lattice state is poorly recognized by tau. Fourth, MTs stabilized by expanding ligands (PTX, DTX, Epo B) bound tau more slowly and predominantly through patch formation (**Figure 6B; Supp. Movies 9–10**). These expanded lattices required higher tau concentrations and longer incubation times to reach substantial coverage.

Integrated fluorescence measurements were consistent with these spatial observations (**Figure 6C**): compact lattices accumulated tau rapidly, whereas expanded lattices required higher concentrations and longer times to reach comparable signal intensity. Although ZMP-, DDM-, and LAU-stabilized MTs achieved rapid spatial coverage, total fluorescence continued to increase until equilibrium, indicating that uniform coverage does not imply instantaneous maximal occupancy.

A modest correlation between mean MT radius and total tau binding was detected (**Figure 6E** and **6G**). However, longitudinal lattice state was a stronger determinant of tau recognition pattern and binding kinetics than radius alone.

Finally, we performed numerical simulations of a single-step association reaction under conditions in which tau binding proceeds without detectable lattice rearrangement, namely on ZMP-, DDM-, and LAU-stabilized microtubules. Based on the experimentally derived concentrations (5 nM tau; 12.5 nM accessible binding sites), binding can be modeled as a homogeneous one-step interaction (Figure **S3**). Under these conditions, both free tau and free binding sites decrease during the reaction, and the system does not follow strict pseudo-first-order kinetics (48). Simulations reproduce the experimentally observed rapid spatial decoration followed by a slower approach to equilibrium when association rate constants on the order of a few µM^-1^ s^-1^ and dissociation rate constants on the order of 10^−3^ s^-1^ (dissociation half-times of several minutes). A previous washout study of tau dissociation from PTX-stabilized MTs (47) reported half-times on the order of tens of minutes, consistent with the range estimated here. These kinetic estimates represent order-of-magnitude approximations derived from a simplified non-cooperative model and are applicable only to lattice states that do not undergo conformational adaptation during binding. Furthermore, they do not account for cooperative envelope formation or ligand-dependent lattice rearrangements observed in expanded microtubules. Nevertheless, the simulations demonstrate that the measured kinetics are compatible with the high-affinity tau binding and slow dissociation observed under the experimental conditions used.

## Discussion

The discovery of taxanes introduced the possibility of blocking the cytomotive switch in the straight MT conformation (15). By stabilizing polymerized MTs, these compounds suppress controlled force generation during mitosis while preserving polymer mass. This led to the expectation that microtubule-stabilizing agents (MSAs) would cause less neurotoxicity than depolymerizing agents. However, clinical experience revealed that PTX and related taxane-site ligands induce substantial and often persistent neurotoxicity (17, 18), indicating that stabilization itself is not structurally neutral.

Our results (**Figures 1 and 2**) show that MSA-induced stabilization is accompanied by defined lattice remodeling. Rather than creating entirely new architectures, ligands selectively stabilize pre-existing compact (∼4.06 nm) and expanded (∼4.17 nm) longitudinal lattice states. Importantly, lattice expansion is not intrinsically linked to taxane-site occupancy or to stabilization per se. Some taxane-site ligands (e.g., DDM and ZMP) preserve compact lattices, whereas BIII still induces lattice expansion despite exerting only weak functional microtubule stabilization, even at full occupancy. Moreover, small chemical modifications of PTX, such as C-7 substitution, can switch the preferred lattice state. These findings demonstrate that ligand chemistry, rather than binding site alone, determines longitudinal lattice bias.

Although longitudinal spacing exhibits quasi-binary behavior, lateral organization spans a broader structural landscape reflected in shifts of mean radius. Expanded lattices are generally associated with modest decreases in mean radius, whereas compact-stabilizing ligands often produce larger lateral adjustments. However, longitudinal and lateral parameters are partially decoupled: combinations of drugs and nucleotide analogs generate lattices with similar axial spacing but different mean radii. Thus, stabilization and lattice geometry are separable properties.

Time-resolved experiments (**Figure 4**) further clarify the dynamics of these transitions. Longitudinal expansion or compaction occurs within the temporal resolution of the assay, whereas lateral equilibration proceeds more slowly. This asymmetry suggests that the compact and expanded states represent closely spaced conformational minima that can be rapidly interconverted, while redistribution of protofilament organization requires longer equilibration. Together with stoichiometry experiments (**Figure 3**) showing that partial occupancy is sufficient to shift lattice states, these findings support an ensemble-based model in which ligand binding biases a pre-existing equilibrium between conformational states rather than imposing a wholly new structure.

The functional consequences of lattice state selection are substantial (**Figure 7A**). Expanded lattices display reduced apparent GTP hydrolysis rates under steady-state assembly conditions. Molecular simulations indicate that ligand binding subtly reshapes the conformational ensemble of the β1:α2 interdimer interface through helix H7, thereby modulating catalytic-site geometry at the E-site (**Figures S2 and S4**). These effects arise from coordinated, small-amplitude structural adjustments rather than large rearrangements, consistent with the high sensitivity of GTP hydrolysis to precise catalytic alignment (39).

**Figure 7.**
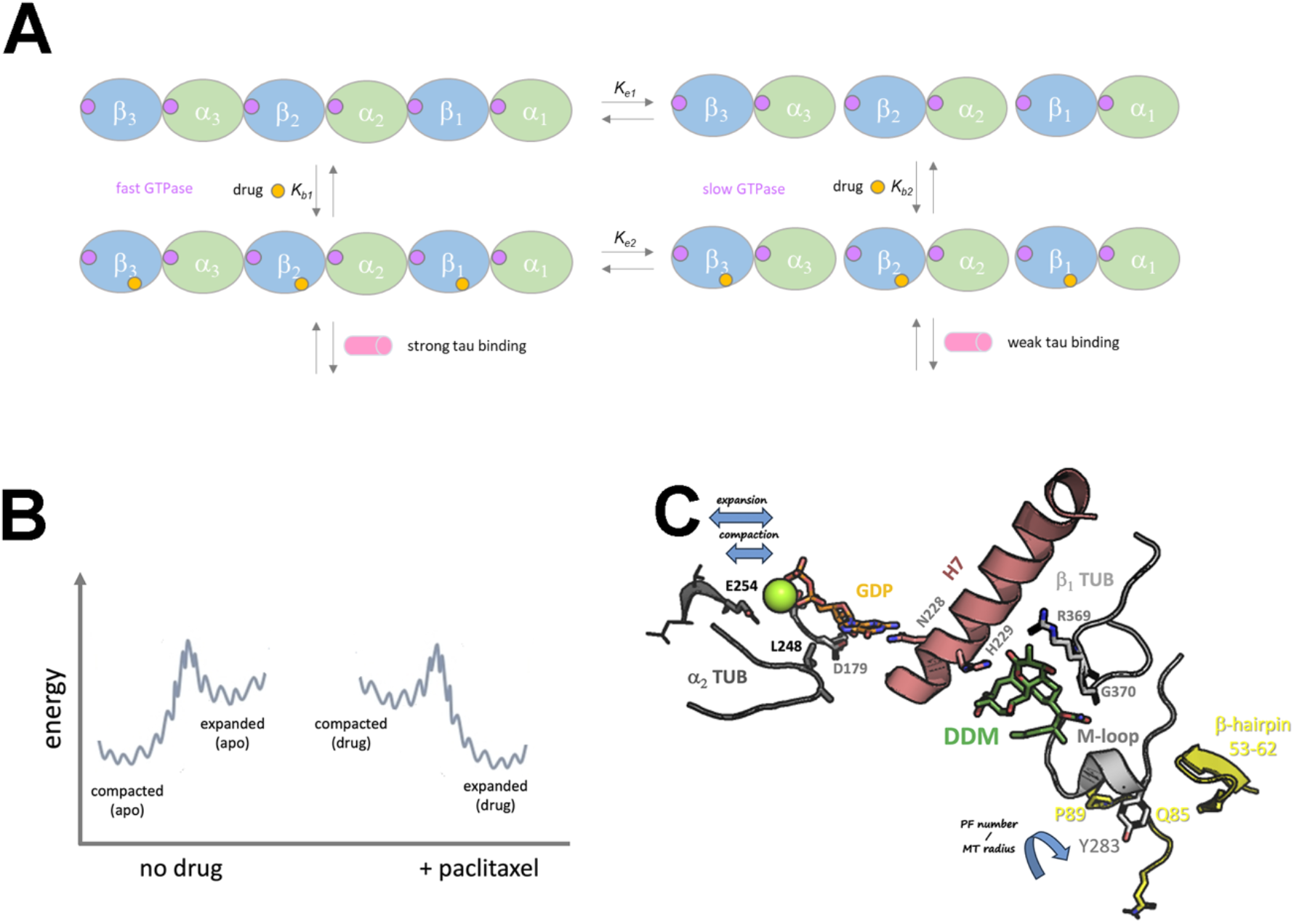
Conceptual model of ligand-induced lattice states and functional modulation. (A) Functional consequences of lattice state selection. Expanded lattices are associated with reduced GTP hydrolysis and weaker tau recognition, whereas compact lattices retain or enhance catalytic activity and promote stronger MAP interaction. The diagram integrates structural state selection with biochemical and regulatory outputs. **(B) Energy landscape of longitudinal lattice states**. Schematic free-energy profile illustrating compact (∼4.06 nm monomer rise) and expanded (∼4.17 nm) lattice conformations as neighboring minima. Ligand binding shifts the population distribution between these states, consistent with the quasi-discrete axial spacings observed experimentally. **(C) Structural coupling between the taxane site and the catalytic E-site**. Simplified cartoon representation depicting the structural basis for the crosstalk at the β1:α2 interdimer interface from the luminal taxane-binding pocket to the E-site through helix H7 of β1-tubulin. Differences in interdimer rise distance and rearrangement of the 178-SDT-180 motif remodel γ-phosphate positioning and water organization around the Mg^2+^ ion (green sphere) for the water-mediated GTP hydrolysis catalyzed by E254 from α2-tubulin. The interprotofilament crosstalk is dominated by the lodging of Y273 into a cavity present in the β1 subunit of the neighboring protofilament (yellow). β1, light gray; α2, dark gray; DDM and GDP are shown as sticks with C atoms colored in green and orange, respectively. The green sphere is the Mg^2+^ ion. ModelArchive DOI: **10.5452/ma-ir5i7**

Beyond catalysis, lattice state also influences MAP recognition and motor behavior. Kinesin velocity is consistently higher on expanded lattices, whereas compressed lattices show greater variability in motor behavior, with broader differences in velocity and run length depending on the ligand present. In particular, strongly stabilizing ligands are associated with shorter run lengths, suggesting that a more rigid lattice may reduce the local structural adaptability required for efficient kinesin stepping. By contrast, lower-affinity stabilizers generally support longer runs. Tau, in contrast, shows a marked preference for compact lattices, binding rapidly and uniformly to some compressed states while interacting more slowly with expanded ones. Notably, tau recognition depends primarily on longitudinal lattice state rather than mean radius alone, reinforcing the central role of axial conformation.

These observations converge in a unified model (**Figure 7**). Compact and expanded lattice states can be conceptualized as neighboring minima along a shared structural landscape (**Figure 7B**). Ligand binding shifts the population distribution between these states without generating entirely new geometries. Structural coupling between the luminal taxane pocket and the interdimer interface provides a pathway through which small chemical perturbations propagate to catalytic and regulatory interfaces (**Figure 7C**). In this framework, lattice conformation encodes functional outputs, linking drug chemistry to enzymatic activity and MAP recognition.

Importantly, structural transitions occur at substoichiometric occupancy and longitudinal states interconvert within seconds, illustrating that modest chemical perturbations are sufficient to bias the conformational ensemble of the polymer. This sensitivity underscores the intrinsic plasticity of tubulin and supports a model in which compact and expanded states are separated by a small energetic difference that can be rapidly shifted by ligand binding. Therefore, MSAs act not only as stabilizers, but as modulators of lattice state.

Here, we have provided a quantitative structural atlas of ligand-induced lattice states and demonstrate that longitudinal conformation functions as a regulatory parameter linking chemical binding to catalytic and MAP-dependent outputs. By showing that ligand chemistry selectively biases rapidly interconvertible lattice states with distinct functional consequences, this work establishes the microtubule lattice as a tunable structural platform and offers a conceptual framework for the rational design of structure-selective modulators with improved therapeutic profiles.

### Summary

Binding of stabilizing drugs to microtubules alters their structure both longitudinally and laterally. Along the longitudinal axis, the lattice adopts two preferred conformational states: a compact state (monomer rise ∼4.06 nm) and an expanded state (∼4.17 nm). These states are closely spaced energetically, as even partial ligand occupancy is sufficient to shift the conformational equilibrium, and longitudinal expansion or compaction occurs within seconds of drug incorporation.

In contrast, lateral organization spans a broader structural landscape reflected in shifts in mean microtubule radius. These variations correspond to differences in average protofilament organization within heterogeneous populations rather than discrete, synchronous protofilament transitions. Although alternative lateral arrangements appear energetically similar, allowing ligand binding to bias their distribution, equilibration toward the final mean radius occurs more gradually than longitudinal changes.

The conformational effects of these compounds can be finely tuned through subtle chemical modifications, such as acylation of the C-7 hydroxyl of PTX. These structural alterations are accompanied by changes in microtubule biochemical properties, including GTPase activity, and significantly influence recognition by tau and kinesin. Together, these findings highlight microtubule architecture as a tunable structural platform that integrates chemical perturbation with catalytic and regulatory outputs.

## Materials and Methods

### Proteins

Purified calf brain tubulin containing a mixture of isotypes (49) was prepared as described (50) and used for biochemical and fiber diffraction experiments. Tau-mCherry was expressed and purified as described previously (41). Human Kif5B–eGFP (amino acids 1–905), containing a C-terminal fluorescent tag followed by a 3C protease cleavage site and a 6×His tag, was expressed and purified as described in (51) with minor modifications. Prior to purification, cells were resuspended in an equal volume of lysis buffer (30 mM HEPES pH 7.5, 300 mM NaCl, 0.5 mM ATP, 5% glycerol, 20 mM imidazole, 0.5 mM Tris(2-carboxyethyl)phosphine (TCEP)) supplemented with 1× protease inhibitor cocktail and benzonase (25 U/mL). Cells were lysed, and the lysate was clarified by centrifugation for 45 min at 40,000 rpm in a 45 Ti fixed-angle rotor (Beckman Coulter) at 4 °C. The supernatant was incubated with pre-equilibrated HisPur™ Ni-NTA agarose resin (Thermo Fisher Scientific) for 1 h at 4 °C with gentle rotation in a gravity-flow column. The resin was washed with lysis buffer and subsequently with buffer B (30 mM HEPES pH 7.5, 300 mM NaCl, 0.5 mM ATP, 5% glycerol, 30 mM imidazole, 0.5 mM TCEP). The protein was eluted after overnight cleavage of the His tag with HRV 3C protease in lysis buffer at 4 °C. The eluate was further purified by size-exclusion chromatography on a Superdex® 200 10/300 GL column equilibrated in exclusion buffer (30 mM HEPES pH 7.5, 300 mM NaCl, 0.5 mM ATP, 5% glycerol, 0.5 mM TCEP). Peak fractions were analyzed by SDS-PAGE, pooled, concentrated using Amicon Ultra-15 centrifugal filter units (Millipore), aliquoted, snap-frozen in liquid nitrogen, and stored at −80 °C.

### Microtubule shear-flow alignment and X-ray fiber diffraction experiments

Fiber diffraction analysis of aligned microtubules was performed as previously described (19), with minor modifications (see **SI**).

Time-resolved X-ray fiber diffraction experiments were performed at the BL40XU beamline of SPring-8 (Hyogo, Japan). A 100 µL aliquot of assembled MTs (100 µM tubulin) was applied to the shear-flow alignment apparatus. At t = 40 s, 1 µL of a 20 mM DMSO stock solution of stabilizing ligand (PTX, DTX, BIII, or LAU) was added by remote-controlled pipette to achieve a final concentration of 200 µM (final DMSO concentration 1% v/v; matched DMSO controls were performed). Diffraction patterns were recorded with a 0.8 m camera length and 0.95 s exposure using a Pilatus 200K detector (Rigaku, Tokyo, Japan). Radial integration and peak extraction were performed using ImageJ (v1.53c, NIH). Spatial calibration was performed using the 4.97 nm powder diffraction signal of lead stearate. One detector pixel corresponded to a reciprocal-space interval of 0.0011 nm^−1^. Peak positions were determined by Lorentzian fitting of the diffraction profiles.

### In silico model building and molecular dynamics simulations

Geometry optimization and derivation of atom-centered RESP charges (52) for DTX, Epo A, ZMP, DDM, and ATX were performed at the B3LYP/6-31G(d,p) level of theory using the IEF-SCRF continuum solvent model (53) for water, as implemented in Gaussian 09 (Revision D.01) (54). This protocol has been applied previously to other tubulin ligands (55) and to covalent adducts of TACCA-AJ and ZMP with Aspβ226 and Hisβ229, respectively (36). Bonded and non-bonded parameters were assigned using the general AMBER force field (GAFF2) (56).

The conformational preferences of DDM in explicit solvent were examined by immersing the ligand (coordinates from PDB entry 6SES) in a cubic box of TIP3P water molecules and simulating at 300 K for 300 ns using molecular dynamics (MD), followed by simulated annealing and energy minimization, as described previously (33, 57).

Macromolecular models representing three parallel short protofilaments (PFs), each consisting of two head-to-tail αβ dimers ((α1:β1–α2:β2)/(α1′:β1′–α2′:β2′)/(α1″:β1″–α2″:β2″)) (MT patches), were constructed from PDB entries 6DPU and 6DPV (58) (expanded and compacted 14-PF MTs, respectively) and 3JAK (59) (symmetrized compact 13-PF MT). Missing residues 38-46 in each α subunit were grafted from the AlphaFold model AF-P81947-F1-v4. Residue numbering and secondary structure assignments follow α-tubulin conventions (60).

Refined 6DPV and 3JAK MT patches were subjected to anisotropic elastic network modeling (ENM) and normal mode analysis (NMA) using the elNémo server (61) with default parameters (12 Å cutoff; 1.0 kcal mol^-1^ Å^-2^ force constant; maximum displacement <2 Å) (62).

The interdimer interface (β1:α2) was further examined using reduced complexes derived from 6DPU (expanded) and 6DPV (compacted). All Cα atoms, except those in flexible loop regions, were restrained using a weak harmonic force constant (2.0 kcal mol^-1^ Å^-2^) to preserve global lattice geometry. GTP at the N-site and either GTP or GDP at the β-tubulin E-site, including partially hydrated Mg^2+^ ions, were positioned according to high-resolution structures (PDB 4I4T and 8BDE). Systems were energy minimized and equilibrated at 300 K for 100 ps using the ff14SB force field (63), GAFF2 for ligands, TIP3P water for explicitly retained water molecules, and the generalized Born model for bulk solvent (64), as implemented in pmemd.cuda_SPFP of AMBER 18. These short simulations were intended for structural relaxation rather than production sampling. Simulated annealing from 300 K to 100 K over 150 ps was subsequently applied.

The different (α1:β1–α2:β2)/(α1′:β1′–α2′:β2′)/(α1″:β1″–α2″:β2″) complexes in apo and holo states were simulated under identical conditions for 200 ps to ensure local equilibration. Ligands were positioned based on experimental binding modes: ATX and DTX were superimposed onto PTX-bound structures (15); DDM and Epo B were docked to reproduce poses observed in PDB entries 6SES and 7DAE; covalent adducts of TACCA-AJ and ZMP were constructed as described (36). Six replicates of each ligand-bound system were sampled in parallel.

The equilibrated MT patches were cooled from 300 K to 100 K over 400 ps and energy minimized without positional restraints. Tubular assemblies were constructed by iterative best-fit superposition of MT patches in PyMOL, following the procedure described in (33). The (α1:β1– α2:β2–α3:β3) stretch of the first copy was superimposed onto the corresponding stretch of the optimized model, and successive copies were iteratively aligned to generate 13- or 14-PF straight microtubule segments (**Figure S5**) with one ligand bound per β subunit. Redundant overlapping coordinates were removed, and the final all-atom models were energy refined and renumbered. Twelve- and fifteen-PF assemblies were generated by superimposing PF segments onto the harmonically deformed geometries obtained from NMA (**Supp. Movie 11**).

Representative models of one complete MT turn with bound PTX, DTX, Epo B, DDM, ZMP, and TACCA-AJ have been deposited in the ModelArchive (www.modelarchive.org) with accession codes ma-rkctk, ma-x9q02, ma-2embb, ma-ir5i7, ma-qcuyh, and ma-qzvr1, respectively. The model containing bound ATX can be inferred from the Flutax-2-bound model (ma-ayvpr) (33). The first non-trivial normal mode calculated for the Cα trace of the refined PTX-bound 12-PF MT illustrates the cooperative closing motion associated with seam sealing (**Supp. Movie 2**).

### Measurement of nucleotide hydrolysis

Nucleotide hydrolysis by MTs was measured in MEDTA buffer (100 mM potassium 2-(N-morpholino)ethanesulfonic acid (K-MES), 1 mM EDTA, 3.4 M glycerol (∼30% v/v), pH 6.7) supplemented with 6 mM MgCl_2_ and 1 mM GTP. Phosphate production was quantified under steady-state assembly conditions using the malachite green inorganic phosphate assay (65). GTPase rates were determined from the linear regime of phosphate accumulation as described in the Supplementary Information.

### In vitro tau and kinesin binding assays on drug-stabilized microtubules

*In vitro* binding of tau and kinesin to drug-stabilized microtubules was performed as previously described (33), with minor modifications (see **SI**).

### Image analysis

For representative figures, data were collected from at least two independent experimental replicates. Unless otherwise stated, image analysis was performed manually using Fiji (ImageJ) (66) (see **SI** for a full description). Graphs and statistical analyses were generated using Sigmaplot 16.0. Data are presented as mean ± standard error of the mean (SEM), unless otherwise specified.

### Disclosure of delegation to Generative AI

The authors declare the use of generative AI in the writing process. The following tasks were delegated to GAI tools under full human supervision: Proofreading and editing, summarizing text. The GAI tool used was: Chat-GPT 5.4. Responsibility for the final manuscript lies entirely with the authors. GAI tools are not listed as authors and do not bear responsibility for the final outcomes.

## Acknowledgements

This work was supported by the Ministerio de Ciencia e Innovación (MICIN) and FEDER (PID2022-136765OB-I00 to J.F.D. and PID2021-123399OB-I00 to M.A.O.), by the European Molecular Biology Organization (EMBO) through a Short-Term Fellowship (No. 11234 to R.P.O.), and by MOSBRI (TNA) under project number MOSBRI-2024-296 to R.P.O. We thank the staff of beamline BL11-NCD-SWEET at the ALBA Synchrotron (Cerdanyola del Vallès, Spain) for excellent technical support. Time-resolved kinetic experiments were performed with the approval of the SPring-8 Proposal Review Committee (proposal numbers 2016B1182, 2020A1332, and 2021A1430). We also gratefully acknowledge Ganadería Fernando Díaz for the supply of calf brains.

