## Supplemental Material for "An atlas of microtubule lattice parameters regulated through ligand binding to the microtubule stabilizing sites"

[1]Unidad BICS, Centro de Investigaciones Biológicas Margarita Salas, Consejo Superior de Investigaciones Científicas, 28040 Madrid, Spain.

[2]Instituto Madrileño de Estudios Avanzados en Nanociencia (IMDEA-Nanociencia), Madrid, Spain.

[3]Laboratory of Biomolecular Research, Department of Biology and Chemistry, Paul Scherrer Institute, Villigen, Switzerland.

[4]Institute of Biotechnology Czech Academy of Sciences, BIOCEV, Prague West, Czech Republic.

[5]Department of Biological Sciences, Graduate School of Science, Osaka University, Japan.

[6]Department of Life Sciences, Faculty of Biological Resources, Prefectural University of Hiroshima, Japan.

[7]Research & Utilization Division, Japan Synchrotron Radiation Research Institute (JASRI), SPring-8, Japan.

[8]ALBA Synchrotron, CELLS, Cerdanyola del Vallès 08290, Spain.

[9]Ferrier Research Institute, and Centre for Biodiscovery, Victoria University of Wellington, PO Box 600, Lower Hutt 5010, New Zealand.

[10]School of Chemical & Physical Sciences, and Centre for Biodiscovery, Victoria University of Wellington, PO Box 600, Wellington 6140, New Zealand.

[11]Department of Chemistry, 1102 Natural Sciences II, and Department of Pharmaceutical Sciences, 856 Health Sciences Road, Suite 5400, University of California, Irvine, CA 92697, USA.

[12]ETH Zurich, Department of Chemistry and Applied Biosciences, Institute of Pharmaceutical Sciences, 8093 Zürich, Switzerland.

[13]Department of Organic and Macromolecular Chemistry, Faculty of Sciences, Ghent University, Krijgslaan 291 (S.4), B-9000 GHENT, Belgium.

[14]Institute of Materia Medica, Chinese Academy of Medical Sciences & Peking Union Medical College, 2A Nan Wei Road, Beijing 100050.

[15]Department of Biomedical Sciences, University of Alcalá, Madrid, Spain.

[16]Department of Biological Sciences, Faculty of Science and Engineering, Chuo University, Tokyo, Japan.

¤These authors have contributed equally.

*Corresponding Author

### SUPPLEMENTARY INFORMATION

#### Supplementary Methods

***Chemicals***

GTP and GMPCPP were purchased from Jena Biosciences (Jena, Germany). Sodium molybdate and malachite green oxalate were from Sigma-Aldrich. PTX was from Alfa Aesar; DTX was kindly provided by Rhône-Poulenc Rorer (Aventis, Schiltigheim, France); CBTX and BIII were from Sigma; Epo A, B, and D, TACCA-AJ, and IBP were from MedChemExpress; LAU and PLA were provided by Victoria University of Wellington. CTX40, 101, and 102; 2-AB-PT; 3′N-AB-PT; FTX-1; FTX-2; FCH-3; Azido-BIII; HXFTX; ATX; DDM; ZMP; DCT; TB282; CSTR; and CW235 were synthesized as described (1-14). FB44 and FB53 were synthesized as described (15). GS305 was synthesized as described (10.26434/chemrxiv-2024-nwb48). The synthesis and characterization of PPH are described in patent PCT/EP2014/075903.

All compounds were dissolved in DMSO to a final concentration of 20 mM, aliquoted, and stored at −80 °C until use.

***Microtubule shear-flow alignment and X-ray fiber diffraction experiments.***

Microtubules were assembled from purified bovine brain tubulin (100 µM) in PM buffer (80 mM K-PIPES, 1 mM EGTA, 1 mM dithiothreitol, 0.2 mM Tris, 3 mM $MgCl_2$, pH 6.8) in the presence of either 2 mM GTP or 0.5 mM GMPCPP. Where indicated, drugs were included during polymerization at the specified concentrations (typically 200 µM for static experiments). Additional nucleotide analog conditions included 2 mM GTP + 10 mM $BeF_3^-$ and 2 mM GTP + 500 µM $AlCl_3$ + 2 mM $KHF_2$ (used as the fluoride source for $AlF_4^-$ formation), as well as combinations thereof in the presence or absence of 200 µM PTX.

Polymerized tubulin was quantified by ultracentrifugation as described previously (16). Under the conditions used, polymer levels ranged from ~85 µM in the absence of drug to ~99 µM in the presence of GMPCPP and DDM. Drug occupancy was quantified by HPLC as described (17). Stoichiometric ratios reported in **Figure 3** were corrected according to measured polymerized tubulin and bound drug concentrations.

Static X-ray fiber diffraction data were collected at the BL11-NDC-SWEET beamline of the ALBA Synchrotron (Cerdanyola del Vallès, Spain) at λ = 0.827 nm as described in (18). Diffraction data were processed using our in-house software FiDAT. Longitudinal lattice spacing (monomer rise) was determined directly from the position of meridional layer lines corresponding to the fourth harmonic of the 4 nm axial repeat, as described (18, 19). This axial parameter is obtained directly from peak positions and does not rely on geometric modeling. Radial structural parameters (mean microtubule radius) were determined from the position of the $J_{02}$ equatorial maximum according to $J_{02} = 1.045/R$ $nm^{-1}$(where R is the mean radius in nm). Protofilament number values reported in **Table I** and **Table SI** are model-derived estimates calculated from the mean radius assuming an average inter-protofilament spacing of 5.7 nm (20) and are provided for comparative purposes only. Errors correspond to the standard error of the mean radius and monomer rise derived from peak-position fitting.

***Measurement of nucleotide hydrolysis.***

Nucleotide hydrolysis by microtubules (MTs) was measured in MEDTA buffer (100 mM potassium 2-(N-morpholino)ethanesulfonic acid (K-MES), 1 mM EDTA, 3.4 M glycerol (~30% v/v), pH 6.7) supplemented with 6 mM $MgCl_2$ and 1 mM GTP. Tubulin was desalted to remove residual phosphate and exchange buffer components, and clarified by ultracentrifugation prior to use. Tubulin concentration was determined spectrophotometrically using $\varepsilon_{(275)}$= 107,000 $M^{-1}$ $cm^{-1}$ and diluted to 25 μM for use.

Polymerization was initiated by raising the temperature to 37 °C in the presence or absence of the indicated ligands. Under these conditions, 85–96% of tubulin was incorporated into microtubules, as determined by sedimentation assays (16). Drugs were present throughout both assembly and subsequent measurements. Microtubule assembly was monitored turbidimetrically at 355 nm.

Phosphate production was quantified using the malachite green inorganic phosphate assay (21). At defined time points (**Figure S1** lower panels), 20 µL aliquots were withdrawn and immediately quenched into acidic malachite green reagent (final 0.7 M HCl), which denatures the protein and releases all inorganic phosphate present at the time of sampling. Absorbance at 650 nm was measured after color development, and phosphate concentrations were determined using sodium phosphate standards.

Because reactions were quenched under strongly denaturing conditions, the assay reports cumulative inorganic phosphate generated from the start of the reaction. To obtain rates, phosphate accumulation was analyzed over time, and only the linear regime of the time course was considered. Early time points corresponding to nucleation and rapid elongation were excluded. The selected linear region corresponds to a phase in which polymer mass has reached steady state and phosphate concentration increases approximately linearly with time (**Figure S1**). The slope of this region, normalized by tubulin concentration, was taken as the apparent GTP hydrolysis rate. Note that under these conditions, the measured rate is affected by steady-state turnover within the polymer and therefore does not fully reflect intrinsic hydrolysis within a static lattice. Nucleotide content in assembled MTs was determined by sedimentation followed by HPLC analysis as described previously (16).

***In vitro tau and kinesin binding assays on drug-stabilized microtubules.***

*In vitro* binding of Kif5B(1–905)-eGFP and tau441-mCherry to drug-stabilized microtubules was measured by total internal reflection fluorescence (TIRF) microscopy as described in (22), replacing PTX with the indicated compounds. TIRF microscopy was performed on an inverted Nikon Ti2-E widefield microscope equipped with a motorized XY stage, Perfect Focus System, Apo TIRF 60× oil objective (NA 1.49), and a PRIME BSY camera (Teledyne Photometrics). Microtubules (MTs) were visualized by interference reflection microscopy (IRM), and fluorescently labeled proteins were visualized sequentially using GFP and mCherry filter cubes. The microscope was controlled using NIS-Elements software (Nikon). All experiments were conducted at room temperature.

Flow chambers were assembled from silanized glass coverslips (22 × 22 mm and 18 × 18 mm) separated by parafilm strips (heated 15–20 s at 60 °C and gently pressed together), generating ~2.5 mm-wide flow channels. Coverslips were washed in 3 M KOH, plasma-cleaned (Zepto Plasma Cleaner, Diener Electronic), and silanized with hexamethyldisilazane. Channels were incubated with 40 µg $mL^{-1}$ anti-biotin antibodies (Sigma, B3640) in PBS for 10 min, followed by blocking with 1% Pluronic F127 (Sigma-Aldrich, P2443) in PBS for at least 1 h. Excess blocker was removed with BRB80 buffer (80 mM PIPES, 1 mM EGTA, 1 mM $MgCl_2$, pH 6.9).

MTs were assembled from porcine brain tubulin purified using the high-molarity PIPES procedure (23). Polymerization reactions contained 2% biotinylated tubulin (Cytoskeleton Inc., T333P). Drug-stabilized MTs were polymerized from 4 mg $mL^{-1}$ tubulin for 30 min at 37 °C in BRB80 supplemented with 4 mM $MgCl_2$, 1 mM GTP, 40 µM of the indicated drug, and 5% DMSO. MTs were pelleted for 30 min at 18,000 × g (Microfuge 18, Beckman Coulter), resuspended, and stored in BRB80 containing 10 µM drug. All drug-stabilized MTs were stored and imaged in the continuous presence of the corresponding ligand. GMPCPP-stabilized MTs were assembled from 4 mg $mL^{-1}$ tubulin for 2 h at 37 °C in BRB80 supplemented with 1 mM $MgCl_2$ and 1 mM GMPCPP. MTs were pelleted at 18,000 × g for 30 min and resuspended in BRB80.

Stabilized MTs were introduced into the flow channel and allowed to attach for around 10 s. Unbound MTs were removed by washing with BRB80 containing 10 µM of the corresponding drug. Prior to imaging, buffer was exchanged to motility buffer (BRB80 containing 10 mM DTT, 20 mM D-glucose, 0.1% Tween 20, 0.5 mg $mL^{-1}$ casein, 1 mM Mg-ATP, 0.02 mg $mL^{-1}$ catalase, 0.22 mg $mL^{-1}$ glucose oxidase, and 10 µM drug). Subsequently, either 1.08 nM Kif5B(1–905)-eGFP or 5/10 nM tau441-mCherry in motility buffer was introduced into the chamber.

For kinesin motility assays, an IRM image (50 ms exposure) was acquired, followed by a 1 min video under 488 nm excitation (20% laser power, 300 ms exposure) with no delay. Three to four videos were recorded from different fields within the same channel, with at least two channels analyzed per condition. Experiments were repeated on 2–3 independent days, yielding a total of 36-48 analyzed MTs per replicate.

For tau binding assays, 4 min videos were acquired at 7 s intervals. IRM (50 ms exposure) and 561 nm fluorescence (30% laser power, 200 ms exposure) were recorded sequentially. Depending on the drug used, firstly 5/10 nM tau-mCherry was injected immediately after the first frame was acquired. Similarly, a second/third video of the same area was recorded adding up to 10/40 (second video) and 40 nM (if third video) tau-mCherry.

Because MTs were immobilized on a surface and non-immobilized ones washed afterward, the effective concentration of accessible binding sites cannot be precisely defined; nevertheless, we can place an upper bound on the effective concentration of binding sites. Since the chamber volume is approximately 8 µL, and in the tau experiments this protein is introduced by flowing ~20 µL of a 5 nM solution and ZMP- stabilized MTs reach saturation under these conditions and do not exhibit increased binding upon further increases in tau concentration, the total accessible binding site concentration must be in the low-nM range, with an upper bound of approximately 12.5 nM. Tau binding kinetics were qualitatively analyzed using a simple single-step binding simulation (https://www.stolaf.edu/depts/chemistry/courses/toolkits/126/js/kinetics/) using initial concentrations in the low nM range (**Supp. Figure 3;** 12.5 nM binding sites and 5 nM tau). These simulations reproduce both the observed timescale and saturation behavior.

***Image analysis****.*

For representative figures, data were collected from at least two independent experimental replicates. Unless otherwise stated, image analysis was performed manually using Fiji (ImageJ) (24). Graphs and statistical analyses were generated using GraphPad Prism v8.0.1. Data are presented as mean ± standard error of the mean (SEM), unless otherwise specified.

Background fluorescence was subtracted from all intensity measurements prior to analysis. For each condition, 12 MTs per channel were randomly selected among well-separated, in-focus filaments identified in the IRM channel. Selection was performed prior to fluorescence quantification to avoid bias. MTs exhibiting overlaps, obvious defects, or drifting during acquisition were excluded from analysis.

For kinesin motility measurements, MTs were traced in the IRM channel using segmented lines 5 pixels wide. Kymographs were generated using the *KymographBuilder* plugin in Fiji. Individual motor trajectories were manually traced as 1-pixel-wide straight lines on the kymographs. From these traces: Run length was determined from the displacement along the x-axis. Run duration (dwell time) was determined from the y-axis. Velocity was calculated as run length divided by run duration. Landing rate was calculated by counting the total number of motor binding events per MT and normalizing to both the MT length (µm), acquisition time (s), and kinesin concentration, being reported as events$\cdot s^{-1} \cdot \mu m^{-1}$ of MT$\cdot nM^{-1}$ kinesin. For each condition, analyses included MTs from at least two independent experiments performed on different days.

To quantify the fraction of MT surface covered by tau envelopes, segmented lines (5 pixels wide; corresponding to 0.36 µm under our imaging conditions) were manually drawn along envelope-covered regions or along the full MT length. The total envelope length was divided by the total MT length to yield the percentage of coverage. Coverage analysis was performed at defined time points: 12 s, 1 min, 2 min, and 4 min for the first acquisition (5 or 10 nM tau), and 12 s, 2 min, and 4 min for subsequent tau concentrations. To quantify tau binding per unit length, segmented lines (5 pixels wide) were drawn along the full MT length. The integrated fluorescence density (Relative I) was measured, background-subtracted, and normalized to MT length (µm), yielding fluorescence intensity per µm. All analyses were performed on at least two independent experiments per condition (24 MTs per replica).

**Supplementary Tables.**

| Drug | Average Radius (nm)[1] | Estimated average PF number (model derived)[2] | 4th layer line position ($nm^{-1}$) | Monomer rise (nm) | Lattice longitudinal state[3] |
|---|---|---|---|---|---|
| **Non drug-stabilized MTs** | | | | | |
| **GMPCPP** | 11.83 ± 0.04 | 13.1 | 0.960 ± 0.001 | 4.17 ± 0.01 | E |
| **GDP·$BeF_3^-$** | 11.18 ± 0.01 | 12.3 | 0.979 ± 0.007 | 4.08 ± 0.03 | C |
| **GDP·$AlF_4^-$** | 11.51 ± 0.06 | 12.7 | 0.986 ± 0.001 | 4.05 ± 0.03 | C |
| **Taxanes** | | | | | |
| **PTX+CPP** | 11.38 ± 0.05 | 12.6 | 0.955 ± 0.003 | 4.18 ± 0.02 | E |
| **PTX+$BeF_3^-$** | 11.32 ± 0.01 | 12.5 | 0.994 ± 0.003 | 4.03 ± 0.03 | C |
| **PTX+$AlF_4^-$** | 11.38 ± 0.01 | 12.6 | 0.991 ± 0.003 | 4.04 ± 0.02 | C |
| **DTX+CPP** | 11.86 ± 0.05 | 13.1 | 0.958 ± 0.001 | 4.17 ± 0.01 | E |
| **CBTX+CPP** | 11.8 ± 0.1 | 13.0 | 0.961 ± 0.001 | 4.16 ± 0.02 | E |
| **BIII+CPP** | 10.83 ± 0.02 | 11.9 | 0.958 ± 0.001 | 4.18 ± 0.02 | E |
| **Azido-BIII+CPP** | 10.82 ± 0.03 | 11.9 | 0.960 ± 0.001 | 4.17 ± 0.01 | E |
| **CTX40+CPP** | 11.09 ± 0.02 | 12.2 | 0.958 ± 0.001 | 4.18 ± 0.01 | E |
| **CTX102+CPP** | 11.55 ± 0.08 | 12.7 | 0.958 ± 0.001 | 4.17 ± 0.01 | E |
| **2-AB-PT+CPP** | 11.85 ± 0.08 | 13.1 | 0.960 ± 0.006 | 4.17 ± 0.03 | E |
| **FTX1+CPP** | 12.91 ± 0.03 | 14.2 | 0.960 ± 0.001 | 4.16 ± 0.01 | E |
| **FTX2+CPP** | 12.97 ± 0.03 | 14.3 | 0.958 ± 0.001 | 4.17 ± 0.01 | E |
| **FB44+CPP** | 12.80 ± 0.05 | 14.1 | 0.958 ± 0.002 | 4.17 ± 0.01 | E |
| **FB53+CPP** | 11.79 ± 0.04 | 13.0 | 0.97 ± 0.01 | 4.12 ± 0.04 | B |
| **HXFTX+CPP** | 13.31 ± 0.02 | 14.7 | 0.963 ± 0.002 | 4.15 ± 0.05 | B |
| **ATX+CPP** | 11.75 ± 0.04 | 13.0 | 0.958 ± 0.001 | 4.18 ± 0.01 | E |
| **Epothilones** | | | | | |
| **Epo A+CPP** | 11.22 ± 0.05 | 12.4 | 0.959 ± 0.01 | 4.17 ± 0.01 | E |
| **Epo B+CPP** | 11.32 ± 0.06 | 12.5 | 0.963 ± 0.002 | 4.16 ± 0.01 | E |
| **Epo D+CPP** | 11.29 ± 0.05 | 12.4 | 0.957 ± 0.004 | 4.18 ± 0.02 | E |
| **IBP+CPP** | 11.4 ± 0.1 | 12.6 | 0.960 ± 0.001 | 4.17 ± 0.01 | E |
| **Zampanolides** | | | | | |
| **ZMP+CPP** | 13.31 ± 0.03 | 14.7 | 0.986 ± 0.001 | 4.06 ± 0.01 | C |
| **DCT+CPP** | 11.86 ± 0.05 | 13.1 | 0.958 ± 0.001 | 4.17 ± 0.01 | E |
| **TB282+CPP** | 13.01 ± 0.02 | 14.4 | 0.97 ± 0.01 | 4.12 ± 0.05 | B |
| **GS305+CPP** | 13.33 ± 0.02 | 14.7 | 0.982 ± 0.007 | 4.07 ± 0.01 | B |
| **Discodermolide** | | | | | |
| **DDM+CPP** | 11.6 ± 0.1 | 12.8 | 0.990 ± 0.003 | 4.04 ± 0.02 | C |
| **Peloruside/Laulimalide** | | | | | |
| **LAU+CPP** | 12.41 ± 0.03 | 13.7 | 0.97 ± 0.01 | 4.12 ± 0.05 | B |
| **PLA+CPP** | 10.65 ± 0.05 | 11.7 | 0.958 ± 0.001 | 4.17 ± 0.01 | E |
| **PPH+CPP** | 11.53 ± 0.02 | 12.7 | 0.959 ± 0.001 | 4.17 ± 0.01 | E |
| **CW235+CPP** | 11.98 ± 0.03 | 13.2 | 0.958 ± 0.001 | 4.17 ± 0.01 | E |

**Table SI. Structural parameters of taxane and peloruside/laulimalide site ligands stabilized microtubules in the presence γ-phosphate analogs or GMPCPP.**

1.- Reported SEMs correspond to the uncertainty in the determination of the mean radius from diffraction measurements, not to the dispersion of protofilament numbers within the heterogeneous microtubule population.

2.- Protofilament number was calculated from the mean microtubule radius assuming an average inter-protofilament distance of 5.7 nm (20). The reported values represent model-derived estimates obtained from geometric conversion of the measured mean radius and are provided for comparative purposes only. Protofilament numbers are therefore rounded to the nearest decimal place.

3.- Lattice longitudinal state classification: C, compressed; E, expanded; B, blurred (heterogeneous).

| Drug | Average Radius (nm)[1] | Estimated average PF number (model derived)[2] | 4th layer line position ($nm^{-1}$) | Monomer rise (nm) | Lattice longitudinal state[3] |
|---|---|---|---|---|---|
| PTX | 11.28 ± 0.03 | 12.4 | 0.955 ± 0.001 | 4.18 ± 0.01 | E |
| DTX | 11.68 ± 0.03 | 12.9 | 0.955 ± 0.005 | 4.18 ± 0.03 | E |
| BIII | 11.02 ± 0.01 | 12.2 | 0.955 ± 0.006 | 4.18 ± 0.03 | E |

**Table SII**. Structural parameters of apo assembled microtubules after 30 min incubation with a 2x excess of taxane site ligands.

1.- Reported SEMs correspond to the uncertainty in the determination of the mean radius from diffraction measurements, not to the dispersion of protofilament numbers within the heterogeneous microtubule population.
2.- Protofilament number was calculated from the mean microtubule radius assuming an average inter-protofilament distance of 5.7 nm (20). The reported values represent model-derived estimates obtained from geometric conversion of the measured mean radius and are provided for comparative purposes only. Protofilament numbers are therefore rounded to the nearest decimal place.
3.- Lattice longitudinal state classification: C, compressed; E, expanded.

***Supplementary Figures.***

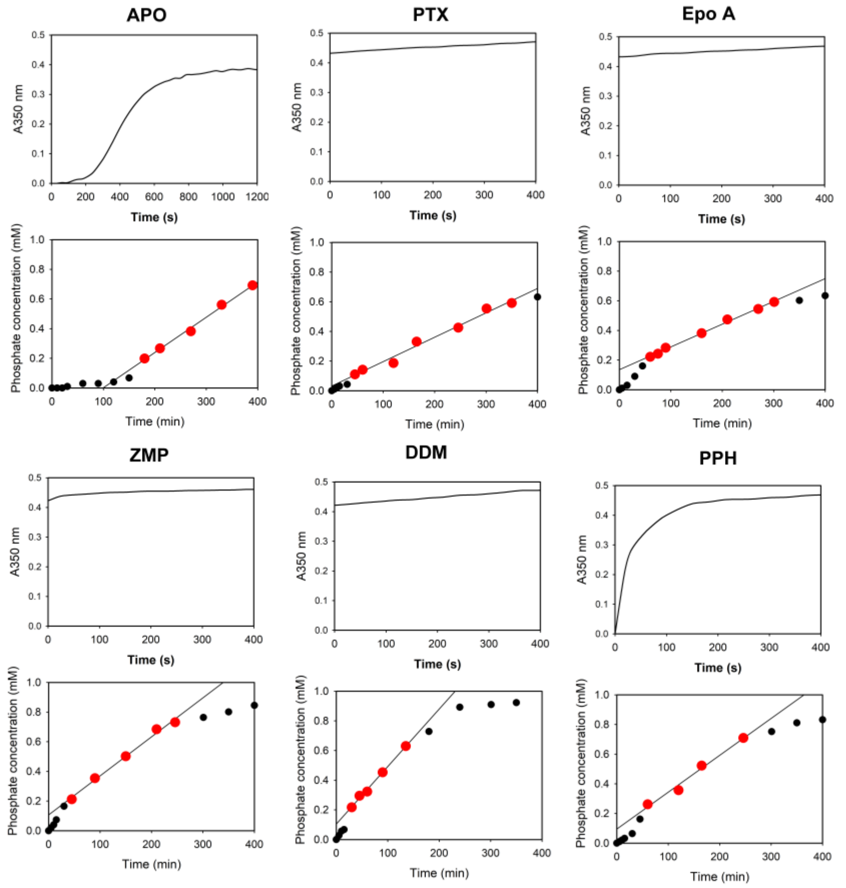


**Figure S1. Representative analysis of microtubule assembly and GTP hydrolysis under stabilizing conditions.** Representative data illustrating microtubule polymerization and associated phosphate production in the absence and presence of stabilizers. Upper panels show microtubule assembly monitored by turbidity at 350 nm, reflecting the kinetics of polymer formation. Lower panels show the corresponding accumulation of inorganic phosphate over time, measured using the malachite green assay. Phosphate production exhibits three phases: an initial lag phase (black small dots), a linear phase characterized by a constant rate of phosphate accumulation (red big dots), and a plateau phase as GTP becomes depleted (black small dots). The apparent GTP hydrolysis rate was determined exclusively from the linear regime (highlighted data points), where phosphate concentration increases linearly with time.

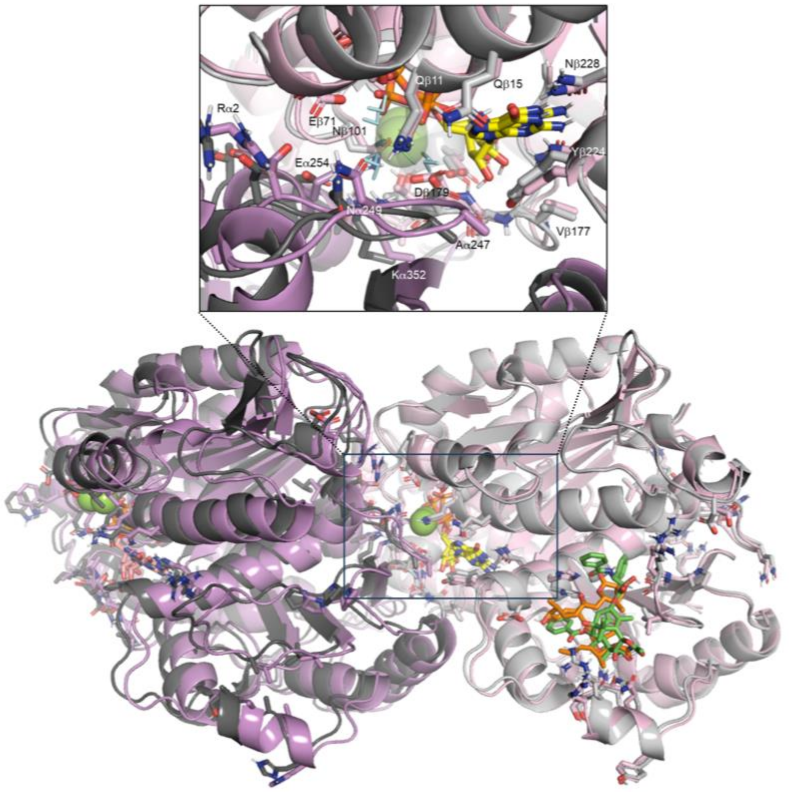


**Figure S2. Structural plasticity of the β1:α2 interdimer interface in compact and expanded lattice states.** Overlay of the β1-α2 interfaces from a compressed PF (light and dark pink) with bound DDM (C atoms in orange) and an expanded PF (light and dark gray) with bound PTX (C atoms in green) using only the β1 subunits for the root-mean-square superposition in PyMOL. The interdimer rise, measured as the distance between the centers of mass of the Cα atoms of β and α subunits, is 40.5 Å for the former and 42.0 Å for the latter. The C atoms of the GDP and GTP nucleotides are colored yellow and salmon, respectively, and the $Mg^{2+}$ ions are displayed as green spheres. The boxed area appears enlarged at the top of figure to show the most notable interfacial differences and the hydrating water molecules (cyan thin sticks). Relevant residues have been labeled.

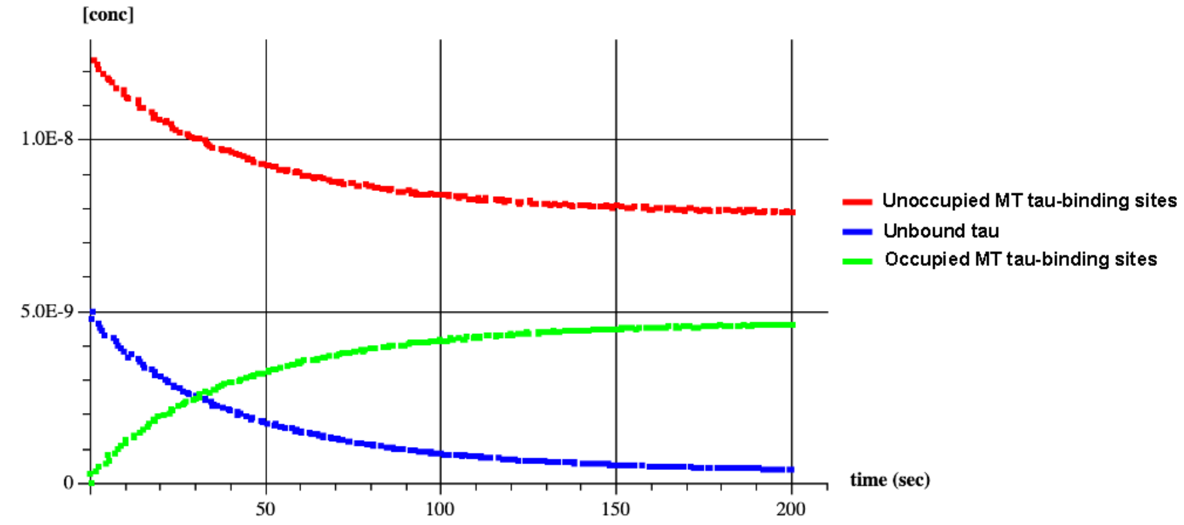


**Figure S3. Simulated binding time course for a single-step association reaction between tau (5 nM) and accessible microtubule binding sites (12.5 nM).** Time evolution of species concentrations obtained from numerical integration of a bimolecular binding model under low nM conditions. Under these conditions, both reactants are progressively depleted, and the system does not follow strict pseudo–first-order kinetics. The selected parameters reproduce the experimentally observed rapid spatial decoration followed by a slower approach to saturation.

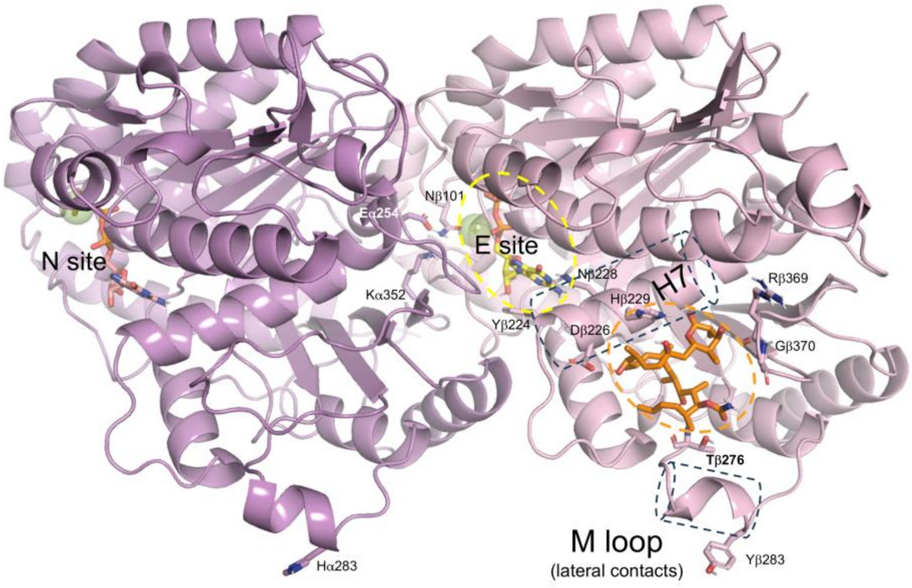


**Figure S4. Helix H7 as a transducer of conformational changes between the taxane-binding site (orange oval), the exchangeable nucleotide site (E site, yellow oval) and the M-loop (an alpha-helix when PFs are associated laterally)**. Ribbon representation of the interdimer interface from a compressed PF (β1, light pink and α2, dark pink) with bound DDM (C atoms in orange) displayed as a representative taxane site-binding ligand. The C atoms of the GDP and GTP nucleotides are colored yellow and salmon, respectively. $Mg^{2+}$ ions are displayed as green spheres and relevant residues mentioned in the text have been labeled.

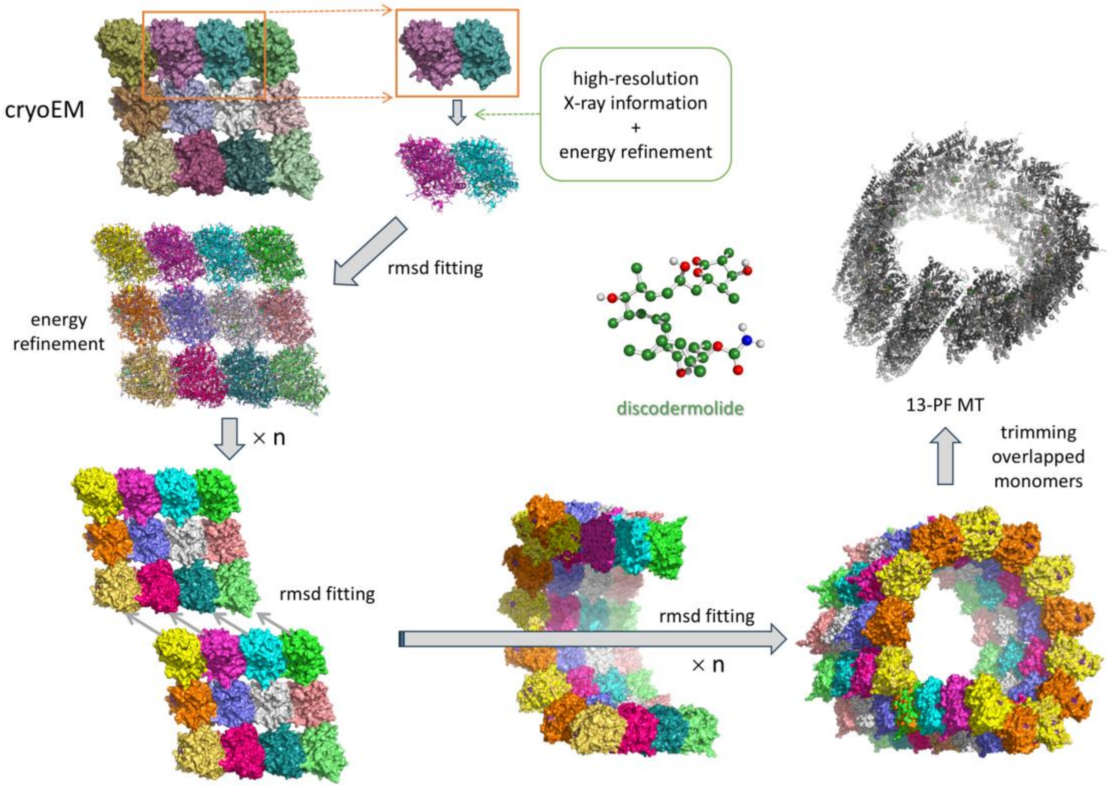


**Figure S5. Schematic describing the procedure followed for constructing a full turn of a 13-PF compacted MT fragment with one discodermolide (DDM) molecule bound to the β subunit of each tubulin dimer** (https://modelarchive.org/doi/10.5452/ma-ir5i7). Briefly, the initial template was provided by a short MT patch consisting of three laterally associated PFs, each made up of two head-to-tail concatenated tubulin dimers, as deposited in the Protein Data Bank with id. 3JAK. The interdimer interfaces in this cryo-EM structure were optimized with the aid of information provided by the high-resolution X-ray crystal structures 8BDE and 8BDF (for the GDP and GTP nucleotides, $Mg^{2+}$ ions and hydration shell waters) and 5LXT (for the ligand bound at the taxane site in β-tubulin) together with energy refinement using the AMBER force field. The resulting optimized patch was replicated six times in PyMOL. The (α1:β1-α2:β2) stretch of the first copy was then best-fit superimposed onto the (α1":β1"-α2":β2") stretch of the original refined model. Thereafter, the (α1:β1-α2:β2) stretch of each remaining copy was best-fit superimposed onto the (α1":β1"-α2":β2") stretch of the preceding model. This process led quite naturally to formation of a tubular structure consisting of 13 straight PFs with one DDM molecule lodged in the taxane site of every β subunit. Lastly, the macromolecular ensemble was (i) cleaned up by removing the redundant subunit copies used for the superposition and (ii) energy refined in the AMBER force field. A similar procedure was followed to obtain the other complexes reported in this paper, as described in the corresponding entries of the ModelArchive repository. For Epo B and ZMP, additional use was made of normal mode analysis to create the 12- and 15-PF MT structures (ModelArchive entries ma-2embb and ma-qcuyh, respectively), as reported previously for paclitaxel (ma-rkctk).

***Supplementary Movies.***

**Supp. Movie 1: Representative time-resolved fiber diffraction images.** Sequence of images showing fiber diffraction patterns of apo-MTs and their structural response to the addition of an 2x excess of drug at time = 40 s. (**A** PTX, **B** DTX, **C** BIII, **D** LAU).

**Supp. Movie 2:** Conformational changes observed by calculating the first non-trivial normal mode for a MT patch consisting of three laterally associated dimers of dimers (PDB id. 6DPV) using the elNemo web server and the default amplitude value. The three views correspond to three orthologous viewpoints. Note that (i) greater amplitudes can be achieved if a bound ligand further stretches the structure along the normal mode and locks the ensemble in this state until ligand dissociation takes place, and (ii) a decrease in lateral curvature will result in an increase in the number of component PFs and vice versa.

**Supp. Movie 3. Representative video of Kif5B-eGFP movement along drug-stabilized MTs.** 1-minute video of 1.08 nM Kif5B-eGFP using a 488 nm laser (20% laser intensity, 300 ms exposure) was acquired with no delay (20 fps).

**Supp. Movie 4. Tau binding to ZMP-stabilized MTs**. 4 min videos of 5 nM tau-mCherry using a 561 nm laser (30% intensity, 200 ms exposure) were acquired every 7 s.

**Supp. Movie 5. Tau binding to DDM-stabilized MTs**. 4 min videos of 5 nM tau-mCherry using a 561 nm laser (30% intensity, 200 ms exposure) were acquired every 7 s.

**Supp. Movie 6. Tau binding to LAU-stabilized MTs**. 4 min videos of 5 nM tau-mCherry using a 561 nm laser (30% intensity, 200 ms exposure) were acquired every 7 s.

**Supp. Movie 7. Tau binding to PLA-stabilized MTs**. Sequential 4 min videos of (**A**) 5 nM, (**B**)10 nM and (**C**) 40 nM tau-mCherry using a 561 nm laser (30% intensity, 200 ms exposure) were acquired every 7 s.

**Supp. Movie 8. tau binding to PPH-stabilized MTs**. Sequential 4 min videos of (**A**) 10 nM and (**B**) 40 nM tau-mCherry using a 561 nm laser (30% intensity, 200 ms exposure) were acquired every 7 s.

**Supp. Movie 9. Tau binding to Epo B-stabilized MTs**. Sequential 4 min videos of (**A**) 10 nM and (**B**) 40 nM tau-mCherry using a 561 nm laser (30% intensity, 200 ms exposure) were acquired every 7 s.

**Supp. Movie 10. Tau binding to DTX-stabilized MTs**. Sequential 4 min videos of (**A**) 10 nM and (**B**) 40 nM tau-mCherry using a 561 nm laser (30% intensity, 200 ms exposure) were acquired every 7 s.

**Supp. Movie 11**: Conformational changes observed by calculating the first non-trivial normal mode for a short full-circle MT consisting of twelve laterally associated PFs using the elNemo web server and twice the default amplitude value. The three views correspond to three orthologous viewpoints. Note that the motion shown will lead to sealing of the PF gap between the first and the twelfth PF.
